# Members of an array of zinc finger proteins specify distinct *Hox* chromatin boundaries

**DOI:** 10.1101/2023.04.25.538167

**Authors:** Havva Ortabozkoyun, Pin-Yao Huang, Edgar Gonzalez-Buendia, Hyein Cho, Sang Y. Kim, Aristotelis Tsirigos, Esteban O. Mazzoni, Danny Reinberg

**Author notes:** These authors contributed equally to this work (Havva Ortabozkoyun, Pin-Yao Huang). (P.Y.H.): Ionis Pharmaceuticals, Carlsbad, CA92010, USA., (H.C.): Developmental Biology Program, Memorial Sloan Kettering Cancer Center, New York, NY10065, USA.

## Abstract

Partitioning of repressive from actively transcribed chromatin in mammalian cells fosters cell-type specific gene expression patterns. While this partitioning is reconstructed during differentiation, the chromatin occupancy of the key insulator, CTCF, is unchanged at the developmentally important *Hox* clusters. Thus, dynamic changes in chromatin boundaries must entail other activities. Given its requirement for chromatin loop formation, we examined cohesin-based chromatin occupancy without known insulators, CTCF and MAZ, and identified a family of zinc finger proteins (ZNFs), some of which exhibit tissue-specific expression. Two such ZNFs foster chromatin boundaries at the *Hox* clusters that are distinct from each other and from MAZ. PATZ1 was critical to the thoracolumbar boundary in differentiating motor neurons and mouse skeleton, while ZNF263 contributed to cervicothoracic boundaries. We propose that these insulating activities act with cohesin, alone or combinatorially, with or without CTCF, to implement precise positional identity and cell fate during development.

## INTRODUCTION

Key to the appropriate development of multicellular organisms is the spatial and temporal regulation of the transcriptome. The dynamic changes to the three-dimensional (3D) spatial organization of the mammalian genome leads to the compartmentalization of genomic regions that are actively transcribed or repressed during the differentiation process^1,2^. Chromatin insulators have emerged as one of the main components of spatial genome organization^3,4^. Although multiple insulator proteins exist in *Drosophila*^4–6^, CTCF has been the major insulator protein recognized in mammals for decades^7,8^. In vertebrate genomes, CTCF and the ring-shaped cohesin complex are essential for maintaining genome organization and the loss of either disrupts the structure of topologically associated domains (TAD)^9,10^. Chromatin loops are the building blocks of the 3D genome organization, and their formation arises from the generally accepted model of cohesin-mediated loop extrusion^11,12^. Several reports indicate that such extrusion is ultimately blocked and stabilized when cohesin encounters CTCF proteins bound to two convergently oriented CTCF DNA-binding sites, which form the loop anchors^10,13–15^.

Notably, cohesin lacks specific chromatin site recognition and importantly, chromatin binding by CTCF does not display any dynamic changes reflective of the newly formed chromatin boundaries that arise during differentiation or in different cell types^16^. For example, *Hox* gene expression patterns emanating from the four *Hox* clusters in mice are regulated in a spatiotemporal manner during development and are determinant to the anterior-posterior identity of cells^17^. While CTCF is pivotal to the formation of distinct chromatin boundaries that determine anterior-posterior identity, its occupation at its defined *Hox* chromatin sites does not diverge during the further process of differentiation^18^. These features beg the question as to what regulates the extensive reconstruction of chromatin boundaries that occur during the differentiation process.

Indeed, additional factors do exhibit site-specific functional roles in insulation, gene expression, and spatial genome organization^19,20^. For example, during cervical motor neuron (MN) differentiation and mouse development, MAZ is required together with CTCF for maintaining the integrity of the borders that demarcate antagonistic anterior and posterior chromatin domains in the *Hox* clusters. This process delimits the spread of the transcription machinery from those chromatin domains active in transcription into those domains that must be repressed to ensure the positional identity of cells^20^. Yet, neither MAZ, given its limited DNA binding sites at the *Hox* clusters, nor CTCF, given its invariable occupation at the *Hox* clusters during differentiation, can account for the further physical restructuring of the chromatin boundaries, which is paramount to cellular fate determination. Thus, we hypothesized the existence of other proteins that cooperate with cohesin to program the formation of appropriate chromatin boundaries that reflect the progress of the differentiation process.

To explore the existence of additional insulators, we performed a genomics analysis to follow cohesin re-location in the absence of the known insulators, CTCF^21–23^ and MAZ^19,20^. Using this strategy, we identified DNA motifs corresponding to transcription factors comprising various specific zinc finger proteins (ZNFs), including PATZ1 and ZNF263, among others. These proteins often co-localize on chromatin with the cohesin component RAD21, in a combinatorial manner. Our genomic analyses performed in several cell types coupled with a series of functional analyses of PATZ1 and ZNF263 during differentiation led to the discovery that PATZ1 and ZNF263 function as insulation factors that are critical to the integrity of genomic boundaries and therefore, cellular fate. Importantly, while PATZ1 is important to establish the thoracolumbar boundary during differentiation *in vitro* and mouse development, MAZ establishes the cervicothoracic one at the *Hox* clusters^20^. Notably, ZNF263 contributes to the formation of cervicothoracic boundaries at the *Hox* clusters during differentiation, similar to MAZ. Our findings indicate that the differentiation process entails a set of discrete factors that function in the context of cohesin to establish distinct chromatin boundaries *in vivo*. Subsets of these insulating activities appear to function in combination to generate a given chromatin boundary and strikingly, CTCF is not always integral to the program.

## RESULTS

### Chromatin barrier functions determine cohesin localization

While CTCF functions to block cohesin movement in loop-extrusion models^11,12,24,25^, its degradation in mESCs (ΔCTCF) revealed that MAZ chromatin binding overlaps that of cohesin (Figure 1A-C and S1A-B). To ascertain whether the chromatin localization of cohesin is dependent on proteins other than the CTCF and MAZ insulators and to set the stage for identifying such putative insulators, we examined cohesin chromatin localization in the absence of both CTCF and MAZ. We generated a *Maz* KO of both MAZ isoforms in the CTCF-degron mESCs (Figure 1D). The *Maz* KO maintained its morphological integrity and showed comparable levels of CTCF, RAD21, and OCT4, a marker for ESCs, relative to the WT case (Figure 1D). While the loss of both CTCF and MAZ (ΔCTCF/ΔMAZ) was also ineffectual with respect to RAD21 protein levels (Figure 1D), RAD21 now exhibited a markedly reduced occupancy at its recognized sites of re-localization in ΔCTCF (Figure 1E, and see S1C-D for all clusters). This phenomenon is exemplified by comparing the re-localized RAD21 at *Nat9* and *Dnajb12* gene loci in ΔCTCF versus ΔCTCF/ΔMAZ (Figures 1F-G). Thus, the absence of the CTCF and MAZ insulators abolished the predetermined presence of cohesin within chromatin and redirected cohesin to sites occupied by other, yet to be determined insulating factors.

**Figure 1.**
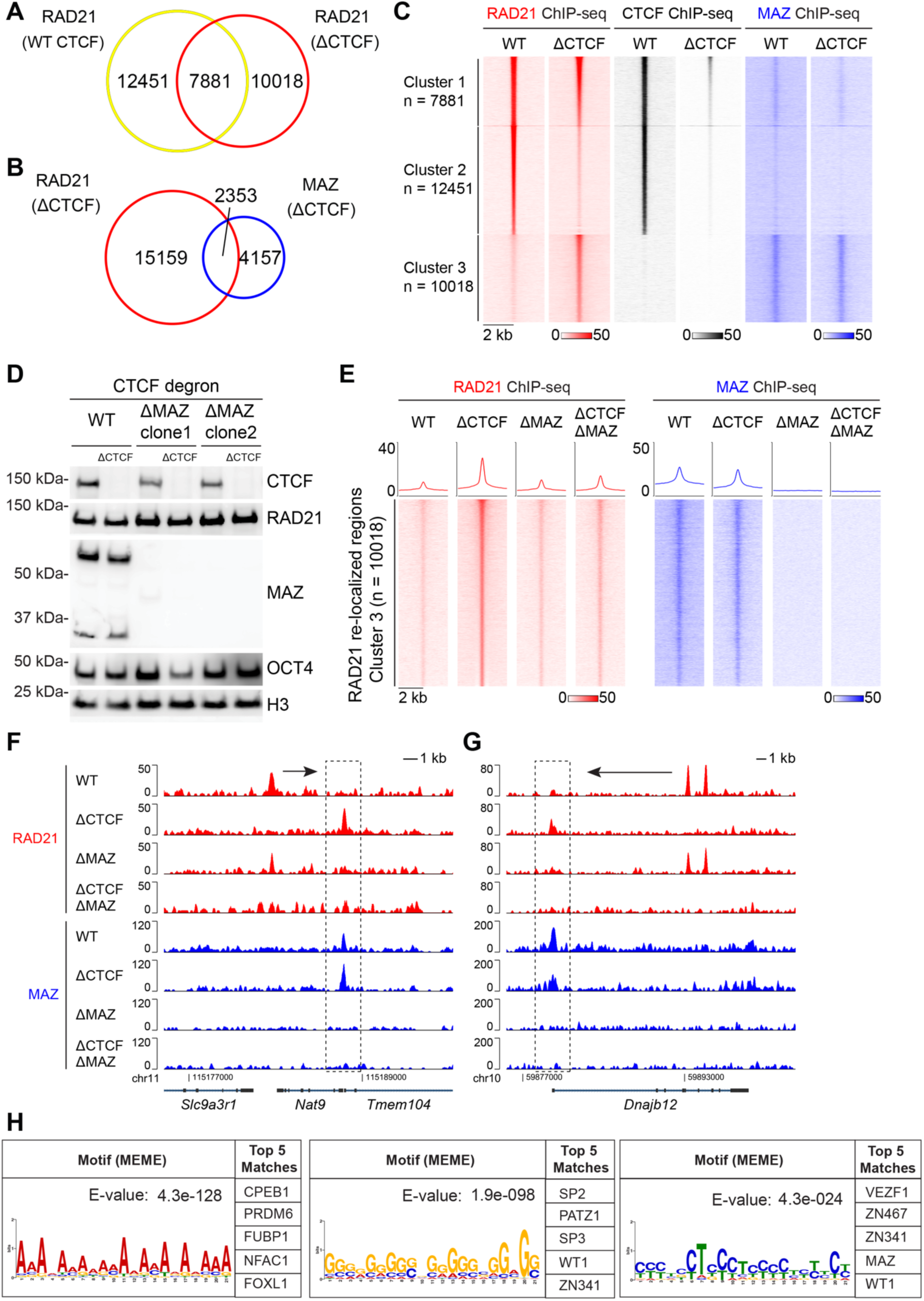
Loss of MAZ in ΔCTCF mESCs reduced RAD21 re-localization, indicating a possible barrier function for MAZ. (A) Venn diagram showing RAD21 binding in mESCs with CTCF intact (WT, Untreated) versus CTCF degraded (ΔCTCF, Auxin treatment, 48 hr). (B) Venn diagram showing MAZ and RAD21 binding in ΔCTCF mESCs. (C) Heat maps of RAD21, CTCF, and MAZ ChIP-seq read density grouped as Cluster 1 (n=7881), Cluster 2 (n=12451), and Cluster 3 (n=10018) based on the alteration in RAD21 signal upon CTCF degradation within a 4 kb window. ChIP-seq data is from one representative of two biological replicates for CTCF and MAZ, and one biological replicate for RAD21. Additional RAD21 ChIP-seq datasets were reported earlier ^15,20^. (D) Western blot analysis of CTCF, MAZ, RAD21, OCT4, and Histone H3 under WT (Untreated), ΔCTCF (Auxin treatment, 48 hr), ΔMAZ, and ΔCTCF/ΔMAZ conditions in ESCs. ΔMAZ represents two independent *Maz* KO clones. (E) Heat maps of RAD21 and MAZ ChIP-seq read density in RAD21 re-localized sites (Cluster 3, n=10018) within a 4 kb window under WT, ΔCTCF, ΔMAZ, and ΔCTCF/ΔMAZ conditions in mESCs (see Figures S1C-D for all clusters). Average density profiles for ChIP-seq under each condition is indicated above the heat map. ChIP-seq data is from one representative of two biological replicates for CTCF and MAZ, and one biological replicate for RAD21. (F-G) Normalized ChIP-seq densities for RAD21 and MAZ at (F) *Nat9* and (G) *Dnajb12* wherein re-localized RAD21 is reduced upon MAZ KO. RAD21-relocalized regions are indicated within dashed-lines. Arrows indicate RAD21 signal in WT that is proximal to the RAD21 re-localized regions. (H) Motif analysis of RAD21 re-localized regions in the absence of CTCF and MAZ by *de novo* MEME motif analysis, along with the corresponding top matches by Tomtom motif comparison (see Figure S1E for the detailed list).

### Uncovering RAD21 re-localization to DNA-binding sites recognized by transcription factors including ZNFs, such as PATZ1

We next sought to uncover possible insulating activities as a function of cohesin chromatin relocalization in the absence of both CTCF and MAZ. Analysis of RAD21 re-localized regions in ΔCTCF/ΔMAZ mESCs revealed the DNA-motifs corresponding to various transcription factors, including the ZNFs (Figures 1H and S1E). Notably, amongst these ZNFs is VEZF1, which was shown previously to function as an insulator^26^, thereby substantiating our strategy for identifying insulation factors.

To gauge the possible co-localization of these ZNFs with RAD21, we analyzed the ChIP-seq data available from public databases (Table S1), in human HEK293, HepG2, and K562 cell lines (Figures 2, and S2-3). Interestingly, amongst the top candidates, PATZ1 (POZ/BTB and AT-Hook-Containing Zinc Finger Protein 1), also known as MAZ-related factor (MAZR), contains six zinc fingers that exhibit a strong homology to those of MAZ and binds *in vitro* to G-rich sequences that resemble the MAZ motif^19,20,27^. Independently of the motif analysis (Figures 1H and S1E), PATZ1 and VEZF1 also appear to be close to MAZ according to the protein sequence analysis of MAZ (Figure S3A).

**Figure 2.**
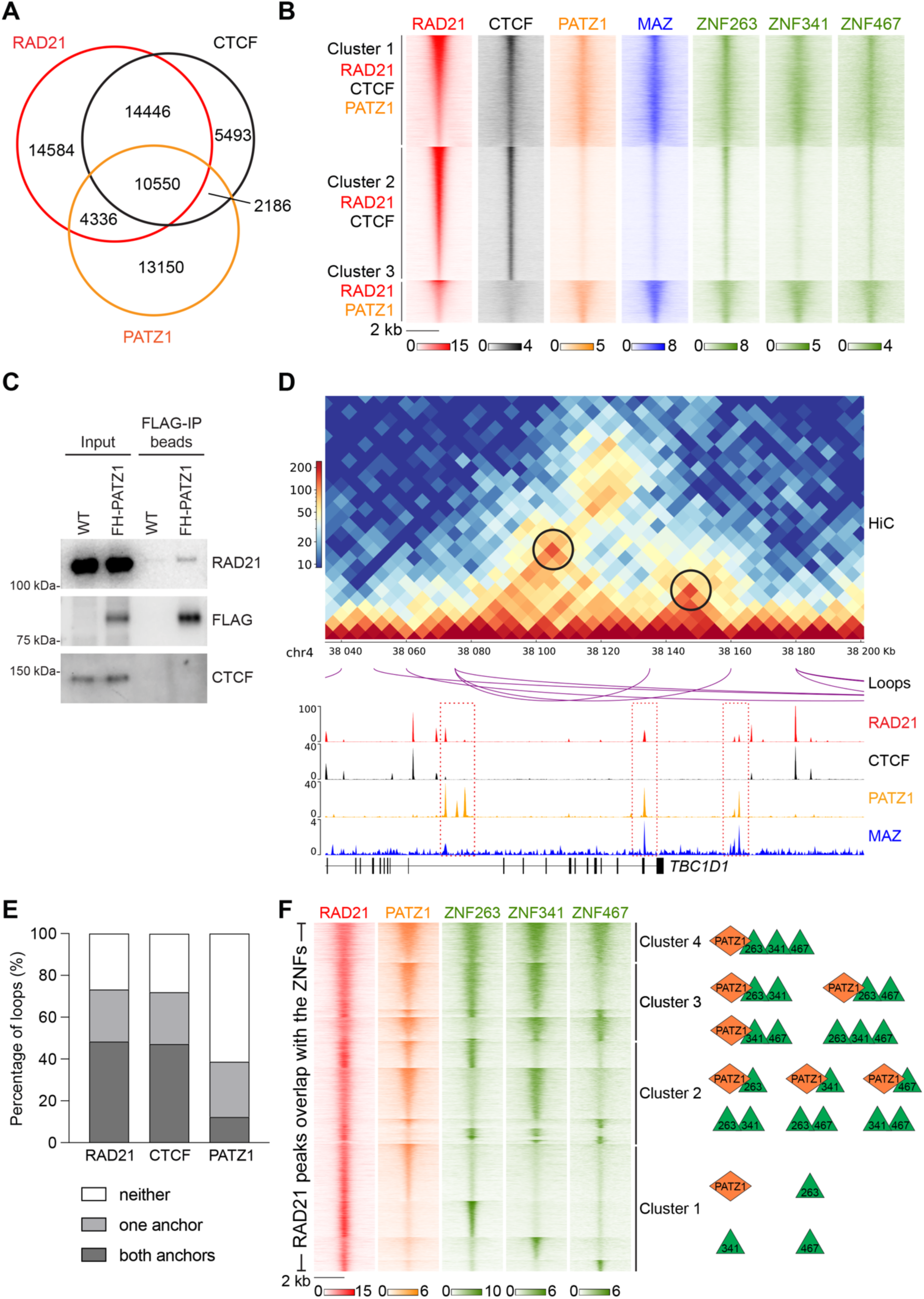
PATZ1 and other zinc finger proteins, ZNF263, ZNF341, and ZNF467, co-localize with RAD21 on chromatin and at loop anchors in HEK293 and HepG2 cells. (A) Venn diagram showing RAD21, CTCF, and PATZ1 binding in HEK293 cells. (B) Heat maps of RAD21, CTCF, PATZ1, MAZ, and other zinc finger proteins, ZNF263, ZNF341 and ZNF467 in HEK293 cells. ChIP-seq read density was grouped as Cluster 1, Cluster 2, and Cluster 3 based on the indicated overlaps with RAD21 signal within a 4 kb window. (C) Western blot analysis of RAD21, FLAG, and CTCF upon FLAG-PATZ1 immunoprecipitation from mESCs (n=2, see Figure S2H for biological replicate). (D) Visualization of Hi-C contact matrices for a zoomed-in region around the *TBC1D1* locus in HepG2 cells. Shown below are loops with PATZ1 at both anchors in HepG2 cells, ChIP-seq read densities for RAD21, CTCF, PATZ1, and MAZ, and gene annotations. ChIP-seq data in HepG2 cells is from two combined biological replicates. (E) Percentage of Hi-C loops in HepG2 cells overlapping with RAD21, CTCF, and PATZ1 ChIP-seq peaks. (F) Heat maps of RAD21, PATZ1, ZNF263, ZNF341, and ZNF467 in HEK293 cells. ChIP-seq read density was grouped as Cluster 1, Cluster 2, Cluster 3, and Cluster 4 based on the combinatorial overlaps of zinc finger proteins with RAD21 within a 4 kb window in HEK293 cells. The model on the right side indicates combinatorial binding of the indicated factors in each cluster (see Figure S4B). ChIP-seq data in HEK293 cells is from one replicate for RAD21 and one representative of two biological replicates for others (see Table S1 for datasets).

### PATZ1 and other ZNFs co-localize with RAD21 on chromatin and at loop anchors

We next examined if PATZ1 functions to anchor chromatin in a manner similar to that of CTCF and of MAZ. We cross-compared the ChIP-seq peaks of RAD21, CTCF, and PATZ1 in HEK293 and HepG2, and clustered the overlapping regions into 3 groups. Cluster 1 consists of 10,550 and 10,262 sites, which are bound by RAD21, CTCF, and PATZ1 in HEK293 and HepG2, respectively (Figures 2A-B, and S2A-B). Cluster 2 represents RAD21 and CTCF co-localized regions, including 14,446 and 31,799 sites in HEK293 and HepG2, respectively (Figures 2A-B, and S2A-B). Interestingly, in Cluster 3, we observed 4,336 and 8,823 sites that are bound by RAD21 and PATZ1 in the absence of CTCF signals, in HEK293 and HepG2, respectively (Figures 2A-B, and S2A-B). These results suggested that indeed, PATZ1 harbors the intrinsic potential of anchoring cohesin on chromatin in different cell types.

We next aligned the ChIP-seq signals of MAZ and other ZNFs, ZNF263, ZNF341, and ZNF467 (see Figure S4A for motifs), based on the aforementioned clusters, using RAD21, CTCF, and PATZ1. Depending on the cluster analyzed, we observed a similar distribution of the signals amongst the ZNFs compared, along with some noteworthy differences (Figure 2B). While Cluster 1 indicates co-localization of ZNFs and MAZ at RAD21, CTCF, and PATZ1 co-occupied regions, Cluster 2 indicates varying signal intensities for ZNFs at RAD21 and CTCF co-occupied regions. Lastly, Cluster 3 shows co-localization of all ZNFs, but not CTCF, at RAD21 and PATZ1 co-occupied regions. For example, ZNF263 and ZNF467 exhibited higher signal intensities in Cluster 2 than PATZ1, MAZ, and ZNF341 and importantly, Cluster 3 displayed less signal intensity for CTCF (Figure 2B), suggesting that each zinc finger protein has distinctive DNA-binding preferences as shown in Figure S4A, although their conserved motifs were described as G-rich sequences (Figures 1H and S1E). At the level of individual loci, we observed that the cohesin peaks co-localized with the ZNF peaks in different combinations and as stated above, sometimes in the absence of CTCF binding (Figures S2C-F). Based on ChIP-seq in HEK293 cells, all clusters shown in Figure 2B appear to be distributed mainly in intronic and intergenic regions (Figure S2G). Altogether, these results support the hypothesis that the ZNF proteins might serve as stable anchors for cohesin, apart from CTCF. Importantly, PATZ1 interacted with RAD21 in mESCs (Figures 2C and S2H), similar to both CTCF and MAZ^19,20,28^, consistent with PATZ1 exerting a possible barrier role in blocking cohesin extrusion. In accordance, analysis of Hi-C data in HepG2 cells revealed that PATZ1 was found at either one or both sides of ∼40% of the loop anchors, and stable loops were detected between PATZ1 binding peaks at regions devoid of CTCF binding (Figures 2D-E). The presence of PATZ1 at CTCF-lacking loop anchors points to the existence of mechanisms fostering cohesin-mediated loop formation. Similar mechanisms may apply to the other ZNFs at different loop anchors in various cell lineages (see below and Discussion).

Among identified candidates, VEZF1 has been one of the factors that binds to the described G-rich sequences (Figures 1H and S1E), and is a previously defined insulator protein^26^ showing protein sequence similarity to MAZ (Figure S3A). For this reason, we further examined VEZF1 binding in comparison to MAZ and PATZ1 in K562 cells. Although VEZF1 co-localizes with MAZ, CTCF and RAD21 in K562 cells, the co-localization between VEZF1 and PATZ1 is rather a smaller fraction of overall VEZF1 bound sites (Figure S3B). Collectively, these results suggest that multiple proteins, including VEZF1, function with RAD21 in establishing chromatin boundaries (see Figure S4A for motifs).

To further evaluate the chromatin binding patterns of PATZ1 and the other ZNF proteins with respect to cohesin anchoring sites, we analyzed the overlap of ChIP-seq peaks of RAD21, PATZ1, ZNF263, ZNF341, and ZNF467 in HEK293 cells (Figure S4B). Overlapping regions were grouped into 4 clusters based on the number of ZNFs co-localized with RAD21 (Figure 2F). Interestingly, the RAD21 peaks co-localized with various combinations of the ZNF protein peaks (Figure 2F), suggesting that the ZNFs might cooperatively or uniquely anchor cohesin on chromatin at different loci and thereby establish locus-specific chromatin boundaries. Moreover, as the ZNFs are distinctively expressed across different tissues and cell types (Figures S4C-E), we speculate that specific cell types utilize unique or varying combinations of these ZNFs to establish cell-specific chromatin boundaries during differentiation and/or development^13,29^.

### PATZ1 co-localizes with RAD21, CTCF, and MAZ on chromatin in mESCs

To further investigate the functional role of the ZNF proteins, we focused on PATZ1 that contains six zinc fingers similar to the MAZ insulator, AT-hook domains, and a POZ domain (Figure 3A). We generated multiple *Patz1* KO lines in mESCs via CRISPR/Cas9 (Figure S5A). As expected, PATZ1 protein signals were lost in the *Patz1* KO lines, while the levels of RAD21, CTCF, and OCT4 were similar to those of WT (Figure S5A), as observed previously for the *Maz* KO^20^. Interestingly, the levels of MAZ were consistently decreased in the *Patz1* KO clones (Figure S5A), suggesting that PATZ1 regulates MAZ expression.

**Figure 3.**
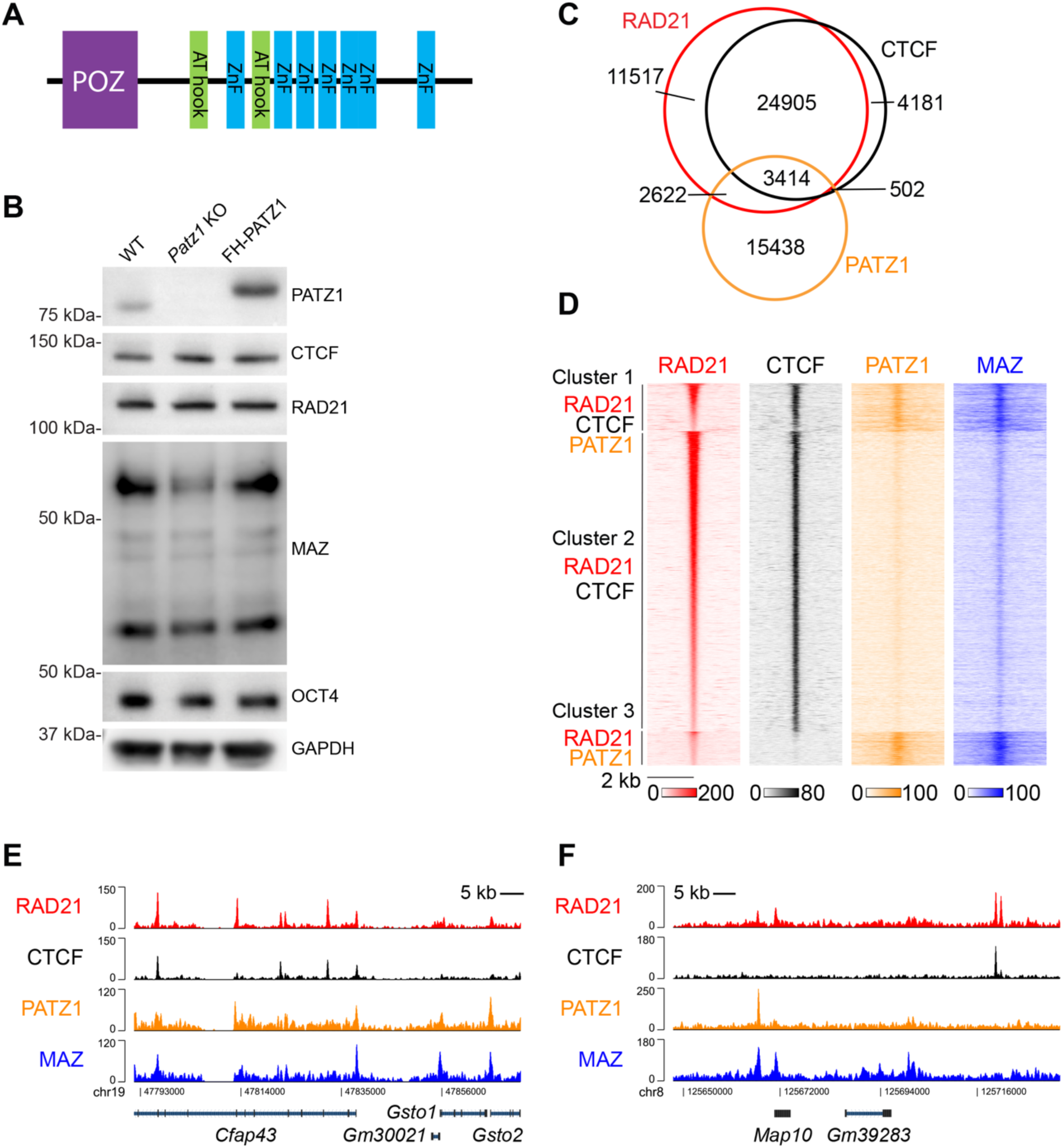
PATZ1 co-localizes with RAD21 on chromatin in mESCs. (A) Schematic of PATZ1 protein domains including zinc fingers (ZnFs), AT hook domains, and the BTB/POZ domain. (B) Western blot analysis of PATZ1, CTCF, RAD21, MAZ, OCT4, and GAPDH in WT, *Patz1* KO, and FH-PATZ1 mESCs (see Figure S5A for four independent *Patz1* KO clones). (C) Venn diagram showing RAD21, CTCF, and PATZ1 binding in mESCs. (D) Heat maps of RAD21, CTCF, PATZ1, and MAZ ChIP-seq read densities grouped as Cluster 1, Cluster 2, and Cluster 3 based on the indicated overlaps with RAD21 signal within a 4 kb window in mESCs. (E-F) Normalized ChIP-seq densities for RAD21, CTCF, PATZ1, and MAZ, wherein co-localizing peaks were visualized. ChIP-seq data is from one representative of two biological replicates.

To determine the genomic binding loci of PATZ1 in mESCs, we introduced 3xFLAG-HA tagged PATZ1 (FH-PATZ1) into the *Patz1* KO cells. The ectopic expression of 3xFLAG-HA-PATZ1 restored both PATZ1 and MAZ levels to the *Patz1* KO, while the levels of RAD21, CTCF, and OCT4 were not altered (Figure 3B). We then determined the genomic binding sites of FH-PATZ1 in mESCs, which gave rise to FH-PATZ1-specific signals as evidenced by the absence of HA ChIP-seq signals in the *Patz1* KO background (Figure S5B). PATZ1 localizes mostly to the promoters, introns, and intergenic regions in mESCs (Figure S5C). Similar to our observations in HEK293 and HepG2 cells, approximately 25% of PATZ1 peaks co-localized with RAD21, either with or without CTCF in mESCs, and MAZ ChIP-seq signals align comparatively with PATZ1 peaks (Figures 3C-D). While PATZ1 containing clusters (Cluster 1 and 3) in Figure 3D are localized mostly to promoters, introns, and intergenic regions, CTCF and RAD21 bound regions (Cluster 2) reside in intronic and intergenic regions (Figure S5D). Indeed, PATZ1 and MAZ form combinatorial binding patterns near RAD21 peaks (Figures 3E-F). Notably, while the binding patterns of PATZ1 and MAZ were similar at the gene loci exemplified in Figure 3E, other loci exhibited differing binding patterns (Figure 3F). These data further support that PATZ1, MAZ, and other ZNFs contribute to cohesin anchoring on chromatin, and that these events are conserved in mouse and human.

### Effects of PATZ1 depletion on cohesin binding and the transcriptome

To further evaluate the molecular function of PATZ1 with respect to cohesin chromatin-binding, we performed ChIP-seq with specific antibodies against RAD21, CTCF, and MAZ in WT and *Patz1* KO mESCs. Approximately 20% of RAD21 peaks and 45% of MAZ peaks exhibited reduced binding upon PATZ1 depletion, while less than 1% of the peaks showed an increase; the reduction in MAZ peaks may be attributable at least in part to the accompanying reduction in MAZ expression in the absence of PATZ1 (Figures S5E-F). On the other hand, 2.6% and 4.5% of CTCF peaks were either increased or decreased, respectively (Figures S5E-F). By comparing the ChIP-seq signals of these factors at the decreased RAD21 peaks, we found that the CTCF signals were minimally altered, whereas MAZ and PATZ1 signals were reduced concomitantly (Figure 4A). Interestingly, inspection of individual loci revealed that the reduction in RAD21 peaks in the absence of PATZ1 could occur in the presence of CTCF binding and irrespective of MAZ binding (Figures 4B-C), suggesting that MAZ and/or PATZ1 facilitates cohesin anchoring.

**Figure 4.**
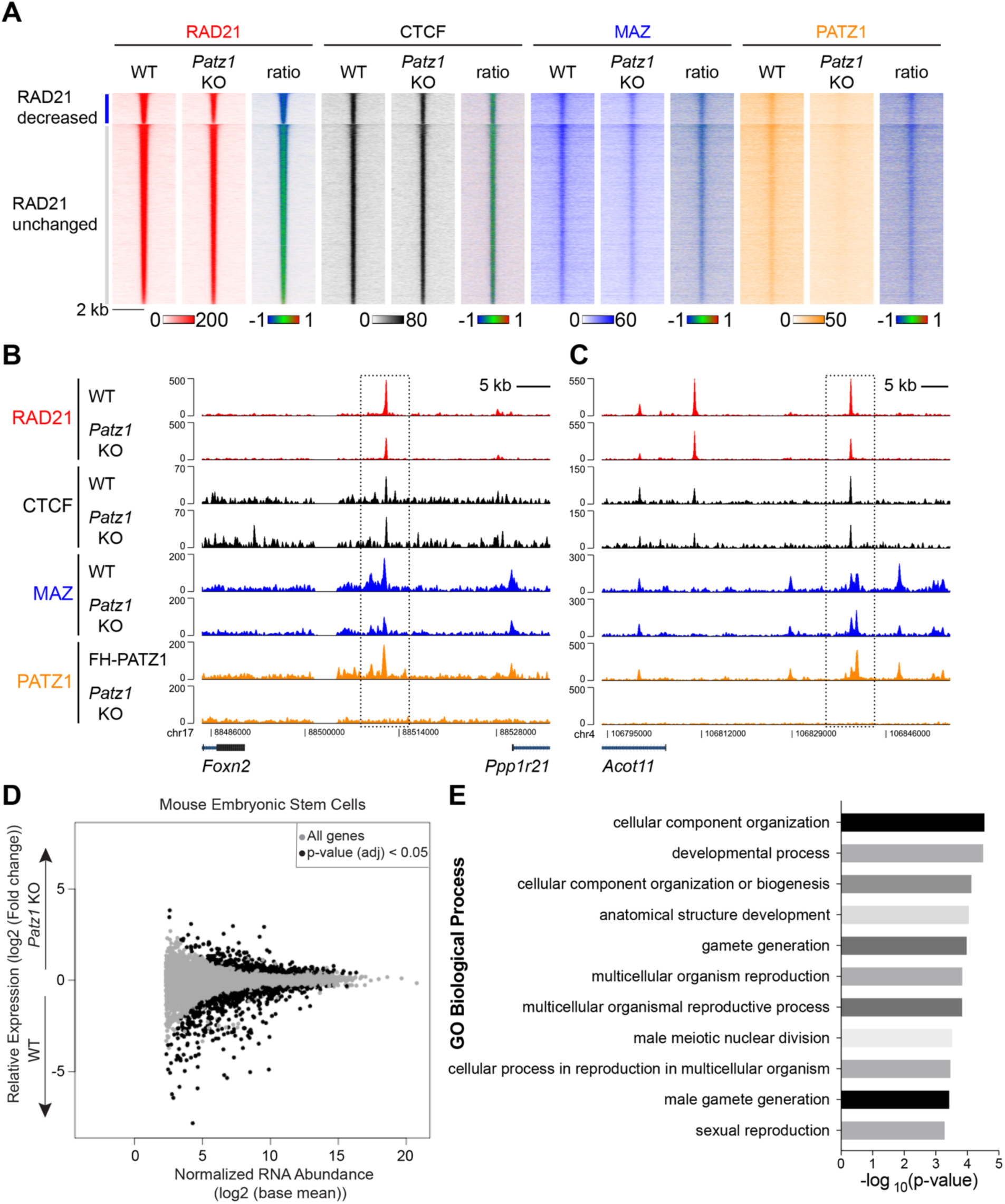
Loss of PATZ1 impacts RAD21 and MAZ chromatin binding in mESCs and gene expression of developmental processes. (A) Heat maps of RAD21, CTCF, MAZ, and PATZ1 ChIP-seq read densities in WT and *Patz1* KO ESCs at RAD21 peaks, clustered into RAD21-decreased and -unchanged sites. Each ratiometric heat map plotted to the right shows the log_2_ fold change (*Patz1* KO/WT) of the signals. (B-C) Normalized ChIP-seq densities for RAD21, CTCF, MAZ, and PATZ1 at the indicated loci wherein MAZ and RAD21 signal was altered. ChIP-seq data is from one representative of two biological replicates. (D) Differentially expressed genes by RNA-seq upon *Patz1* KO in ESCs from three biological replicates (see all in Table S2). (E) Gene Ontology (GO) analysis shows the categories of biological processes that were enriched in the differentially expressed genes in *Patz1* KO versus WT mESCs. PANTHER overrepresentation test tools were used for GO analysis (see all in Table S3).

We next examined the effects of PATZ1 depletion on genome-wide gene expression in mESCs via RNA-seq. The loss of PATZ1 resulted in the differential expression of 1,146 genes (Figure 4D), 585 of which overlapped with PATZ1 ChIP-seq signal (Figure S5G), and 315 of which were similarly impacted in the *Maz* KO (Figure S5H). Importantly, the genes encoding the cohesin loading factors, *Nipbl* and *Mau2*, were expressed at WT levels suggesting that the decreased RAD21 binding observed in the *Patz1* KO was not due to an inherent insufficiency in cohesin loading (Figure S5I). Similar to the *Maz* KO, Gene Ontology (GO) analysis of the *Patz1* KO mESCs revealed an enrichment of several categories related to developmental processes compared to WT (Figure 4E), suggesting that PATZ1 and MAZ might function together to regulate a subset of developmental genes in a cell specific manner.

### Combinatorial binding of insulation factors at *Hox* gene borders is key to proper positional identity in motor neurons

To initiate investigations into how these zinc-finger proteins may function at boundaries, we examined *Hox* gene borders in the context of CTCF, RAD21, and PATZ1 (Figures 5A and S6A-B). As we detected a ChIP-seq peak of PATZ1 at the *Hoxa7|9* border in mESCs, we speculated that PATZ1 may be determinant to the integrity of the *Hoxa7|9* border, as opposed to the requirement of MAZ at the *Hoxa5|6*, as described previously^20^. Thus, we differentiated mESCs into cervical and thoracic MNs using established protocols^30,31^ to foster the formation of a *Hoxa5|6* border and a caudal *Hoxa7|9* border, respectively (Figure S6C). Strikingly, the *Patz1* KO led to de-repression of *Hoxa9* and *Hoxa10* in cervical MNs, as compared to WT (Figure 5B; see Figures 5C-D for other *Hox* clusters). In accordance, the *Patz1* KO appeared to increase *Hoxa7-10* expression in the thoracic MNs, as compared to WT (Figure S6C). The loss of PATZ1 in cervical MNs resulted in the differential regulation of ∼3800 genes (padj < 0.01) with an enrichment of GO categories including cellular division, differentiation, and development processes (Figures 5E-F). Among the differentially expressed genes in PATZ1 KO MNs, 1032 of them (∼27 %) were similarly impacted in MAZ KO (Figure S6D).

**Figure 5.**
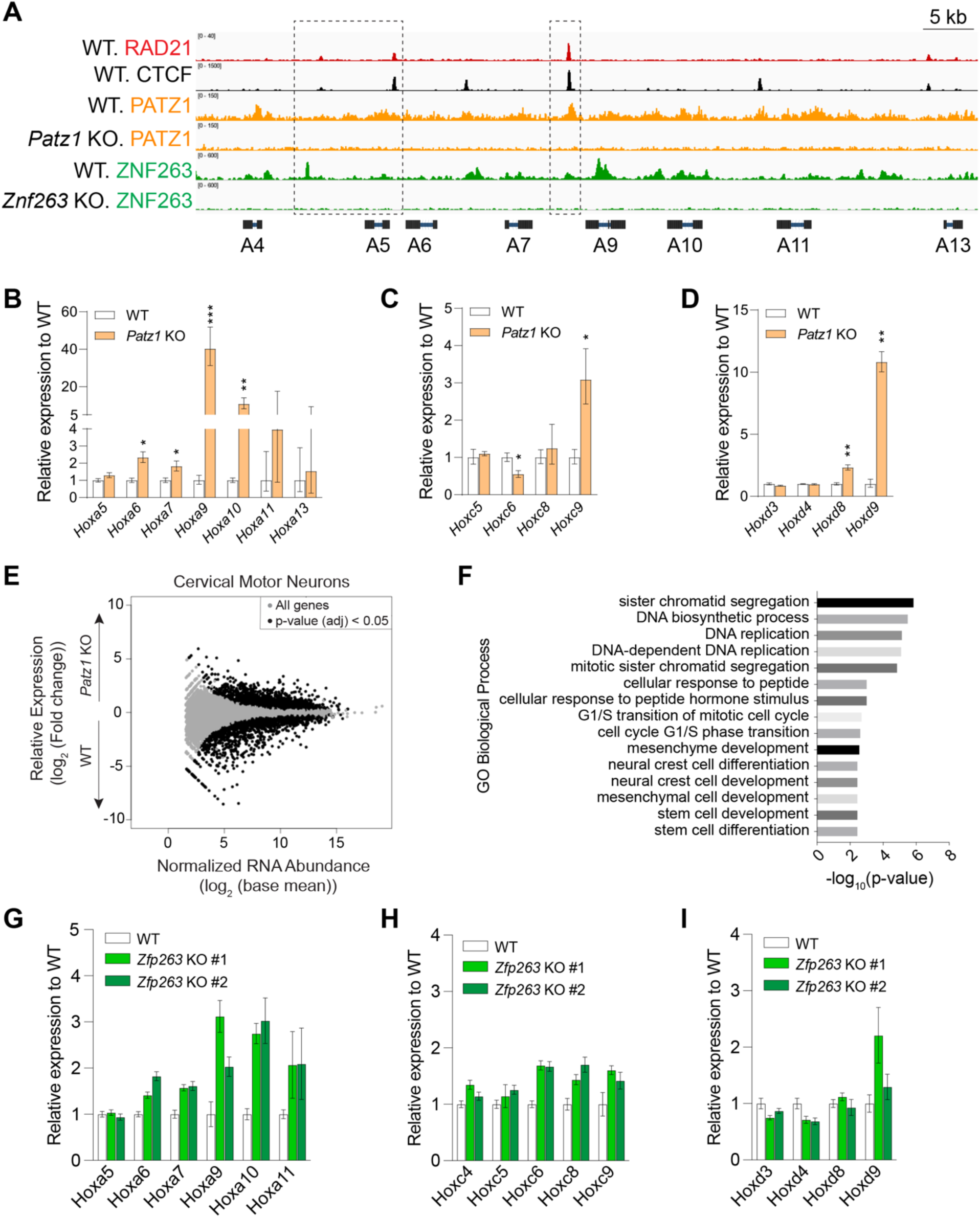
Combinatorial binding of insulation factors at distinct *Hox* gene borders determines rostral-caudal patterning in MNs. (A) Normalized ChIP-seq densities for RAD21, CTCF, and PATZ1, and ZNF263 in WT, *Patz1* KO, and *Znf263* KO mESCs at the indicated regions in the *HoxA* cluster. ChIP-seq data represents one representative replicate of two biological replicates for RAD21, CTCF, and one replicate for FH-PATZ1 and FH-ZNF263. (B-D) RT-qPCR analysis for the indicated *Hox* genes in (B) the *HoxA*, (C) the *HoxC*, and (D) the *HoxD* clusters in WT and *Patz1* KO cervical MNs. RT-qPCR signal was normalized to *Atp5f1* and *ActB* levels. The fold-change in expression was calculated relative to WT MNs. All RT-qPCR results are represented as mean values and error bars indicating log_2_(SE) across three biological replicates (two-sided Student’s *t*-test without multiple testing correction; \*\*\**P* ≤ 0.001, \*\**P* ≤ 0.01, \**P* < 0.05). (E) Differentially expressed genes by RNA-seq upon *Patz1* KO in MNs from two biological replicates (see all in Table S4). (F) GO analysis showing the top biological processes enriched in the differentially expressed genes in *Patz1* KO versus WT MNs. PANTHER overrepresentation test tools were used for GO analysis and top 15 categories having a fold enrichment > 2.5 were plotted (see all in Table S5). (G-I) RT-qPCR analysis for the indicated *Hox* genes in (G) the *HoxA*, (H) the *HoxC*, and (I) the *HoxD* clusters in WT and *Znf263* KO cervical MNs. RT-qPCR signal was normalized to *Atp5f1* and *Gapdh* levels. The fold-change in expression was calculated relative to WT MNs. All RT-qPCR results are represented as mean values and error bars indicating log_2_(SE) across three technical replicates. Two independent *Znf263* KO clones are shown (see Figure S7).

Thus, when the *Patz1* KO and *Maz* KO ESCs were differentiated into cervical MNs, the *Patz1* KO impacted the *Hoxa7|9* border (Figures 5B-D), while the *Maz* KO led to de-repression of earlier *Hox* genes, *i.e.* the *Hoxa5|6* border, as shown previously^20^. Moreover, our analysis of the *HoxC* and *HoxD* clusters pointed to PATZ1 binding preceding the *Hox9* genes (Figures S6A-B), and the loss of PATZ1 led to de-repression of *Hoxc9* and *Hoxd8*-*Hoxd9* (Figures 5C-D), respectively. That the loss of PATZ1 in MNs gave rise to increased *Hoxa9, Hoxc9*, and *Hoxd9* expression, highlights its regulatory role across multiple clusters (Figures 5B-D and S6). As *Hoxc9* upregulation is particularly relevant to the induction of thoracic MN fate^32,33^, *Patz1* KO MNs appear to adopt a more posterior fate during cervical MN differentiation (Figures 5B-D). Importantly, the *Hox* genes retained basal levels of expression in the *Patz1* KO mESCs, similar to that of WT mESCs (Figure S6E), suggesting that PATZ1 does not directly modulate the expression of posterior *Hox* genes, but rather potentiates the establishment of the chromatin boundary upon differentiation.

To investigate whether the loss of other ZNFs identified in this study impacts *Hox* gene borders, we examined *Znf263* KO mESCs during the cervical MN differentiation process (Figures S7A-B). Upon loss of *Znf263,* the levels of RAD21, CTCF, and OCT4 were similar to those of WT in ESCs (Figure S7C). Fittingly, the *Znf263* KO in MNs led to de-repression of *Hoxa6-10* (Figure 5G), *Hoxc6-9* (Figure 5H) and a slight de-repression of *Hoxd9* (Figure 5I; see Figure S7A-B for CRISPR-based genetic deletions). These results suggested that ZNF263 regulates the earlier cervicothoracic border (Figures 5G-I) as opposed to PATZ1 which impacts the thoracolumbar border (Figures 5B-D). In addition, we also note a possible role of ZNF263 in repression of posterior *Hox* genes such as *Hoxa7, Hoxa9* and *Hoxa10* based on the binding pattern in mESCs (Figure 5A). As PATZ1 and ZNF263 co-localize with RAD21 in HEK293 cells (Figure 2), the impact of PATZ1 or ZNF263 loss was analyzed further in MNs. Although about 10-15% of RAD21 binding sites appear to be impacted (Figures S8A-E), we did not observe an alteration of RAD21 binding in the *HoxA* cluster (Figures S8F-G). Thus, both PATZ1 and ZNF263 appear to be critical to the determination of rostro-caudal MN identity and most notably, PATZ1 and ZNF263 regulate distinct *Hox* gene borders. Collectively, the differential roles of PATZ1 and ZNF263 at the *Hox* clusters underscore the existence of a combinatorial code of insulation factors that impact positional and/or cellular identity (see Model in Figures 7A-B).

### Loss of PATZ1 impacts genome organization in ESCs and MNs

In addition to major architectural proteins such as CTCF and cohesin, MAZ contributes to the genome organization. As shown in this study, PATZ1 and several specific ZNFs co-localize with these architectural proteins in multiple cell types examined (Figures 2-5, and S2-3). The loss of PATZ1 impacts RAD21 binding (Figures 4A, S5E-F, and S8) and results in differential expression of 1146 genes in ESCs, and 3800 genes in MNs including *Hox* genes. Thus, we investigated the possible role of PATZ1 on global genome organization by performing Micro-C in WT versus PATZ1 KO ESCs and MNs. As expected, active and repressed compartments had only slight changes between WT and PATZ1 KO (Figures S9A-B). While intra-TAD activities in WT versus PATZ1 KO ESCs were similar, reduced intra-TAD activities were observed in PATZ1 KO MNs compared to WT (Figures S9C-D). Importantly, we observed a downregulation of loops upon loss of PATZ1 in both ESCs and MNs (Figures 6A-D). Aggregate peak analysis (APA) showed a reduction in looping interactions in PATZ1 KO ESCs and MNs compared to WT (Figures 6A-B), indicating the role of PATZ1 in these contacts within the genome. In addition, the differential loop analysis between WT and PATZ1 KO cells indicated alterations in the overall looping interactions (Figures S9E-F). In particular, some gene loci with significant alterations in gene expression (see Tables S2 and S4), reside in regions where loop changes were observed in PATZ1 KO compared to WT (Figures 6C-D and S9G-H). For example, the reduction in looping interactions observed around the *HoxA* cluster in Figure 6C in ESCs correlated with *Hoxa9/10* de-repression and the cell-fate transition phenotype during differentiation (see Figure S8F for cohesin binding). Upon analyzing the common loops between ESCs and MNs, we observed a reduction in looping interactions upon differentiation (Figure S10A-B, top plots). Similarly, PATZ1 loss led to the downregulation of the common loops in ESCs (Figure S10A), while the impact on common loops in MNs was not notable (Figure S10B), suggesting the dynamicity of loop re-organization during differentiation (see Figures 6 and S9). In accordance, PATZ1 was found at one or both sides of ∼35% of loop anchors in ESCs (Figure S10C), as observed in HepG2 cells (Figure 2E). Notably, APA analysis of loops at PATZ1 binding sites showed downregulation in ESCs (Figure S10D), indicating reduced looping interactions at PATZ1 binding regions. Given that the loss of PATZ1 alters the looping interactions between loci and that some of these interactions reflect the concomitant changes in differential gene expression, these findings point to PATZ1 involvement in the three-dimensional genome organization.

**Figure 6.**
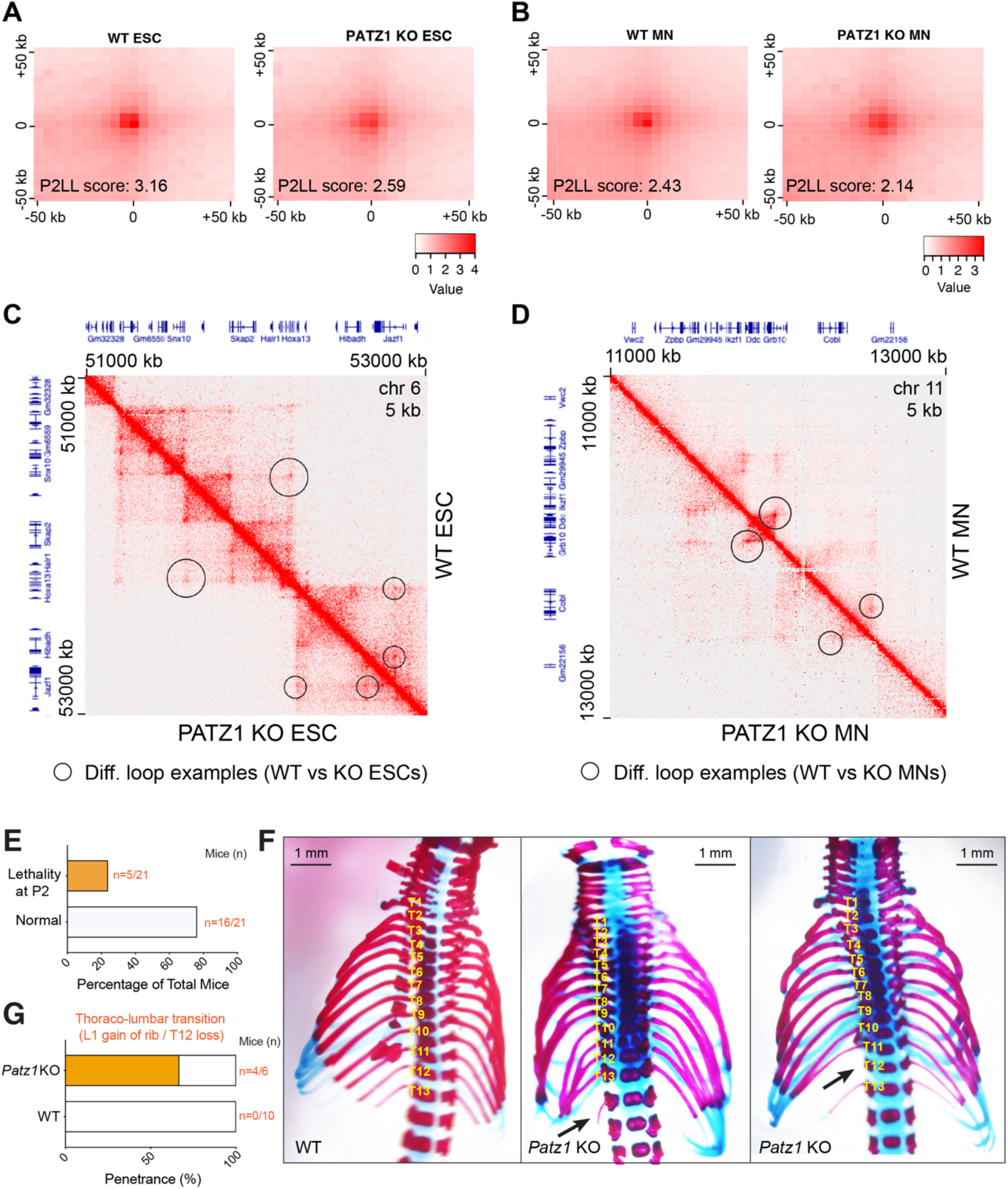
Loss of PATZ1 impacts genome organization *in vitro*, and leads to homeotic transformation at thoracolumbar boundaries in skeletal patterning *in vivo*. (A-B) Normalized APA plots of all loops in WT versus PATZ1 KO (A) ESCs and (B) MNs. The resolution of APA is 5 kb. P2LL (Peak to Lower Left) is the ratio of the central pixel to the mean of the mean of the pixels in the lower left corner. (C-D) Visualization of Micro-C contact matrices for a zoomed-in region around (C) the *HoxA* cluster in WT versus PATZ1 KO ESCs, and (D) *Grb10* locus in WT versus PATZ1 KO MNs. Examples of the differential loops detected have been indicated with the circles. The resolution is 5 kb. Shown above and left are gene annotations. (E) Bar plot showing the percentage of lethality in *Patz1* gRNA injected embryos. The numbers represent pups at postnatal day 2 by which lethality has been observed. (F) Representative Alcian blue–Alizarin red staining of axial skeletons indicating homeotic transformations in WT versus *Patz1* KO mice at postnatal day 0.5. Additional thoracic vertebra, L1 gain of rib, (middle) or loss of T12 (right) are shown with arrows. (G) Bar plot demonstrating the percentage of pups (postnatal day 0.5) with the homeotic transformation phenotype in *Patz1* KO compared to WT. Raw numbers of mice are shown in orange (see Figure S11 for genetic deletions).

### Loss of PATZ1 leads to skeletal patterning defects *in vivo*

As PATZ1 has been found to be critical for proper *Hox* gene expression and positional identity in motor neurons (Figure 5), we hypothesized that the loss of PATZ1 would impact skeletal patterning in mice. Loss of CTCF or MAZ binding were previously shown to impact *Hox* gene boundaries, leading to <u>cervicothoracic</u> transformations of axial skeleton *in vivo*^18,20,34^. Similar to the *Maz* KO^35^, loss of PATZ1 resulted in postnatal lethality within postnatal day 2 (P2) (Figure 6E), as reported earlier^36^, and this outcome was accompanied by skeletal defects at the <u>thoracolumbar</u> boundaries (Figure 6F; see Figure S11 for genetic deletions). The malformations in thoracic vertebra were observed unilaterally with ∼68% penetrance (Figure 6G). These findings strongly support the role of PATZ1 in positional identity determination at the thoracolumbar boundary during skeletal development and highlight the existence of insulation factors functioning in cell fate determination *in vivo* (see Model in Figure 7C).

**Figure 7.**
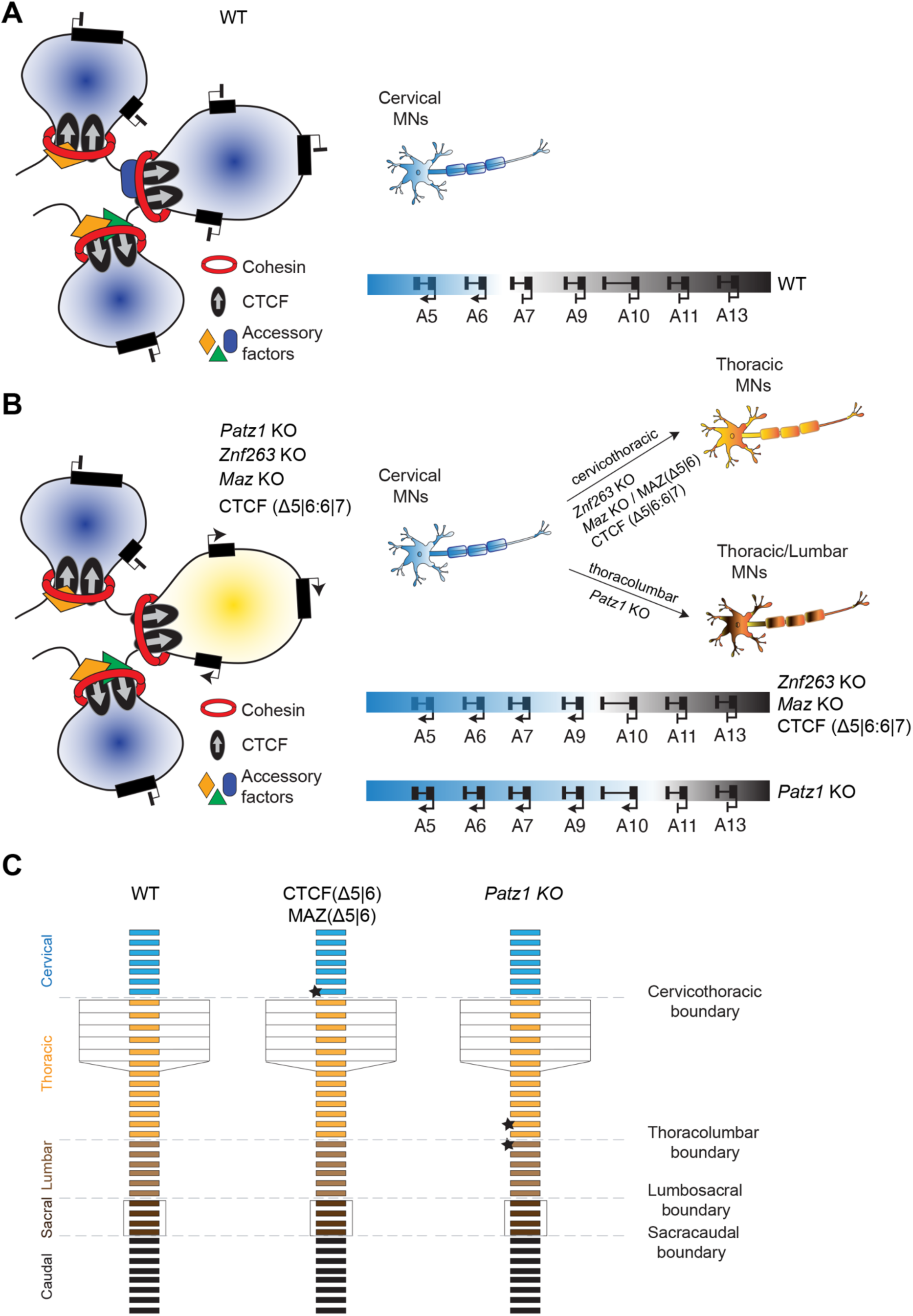
Combinatorial binding of insulation factors at boundaries are critical to the regulation of cellular identity. (A-B) Models depicting (A) the anterior-posterior MN identity in the WT setting in the presence of different accessory/insulation factors and (B) the transition of anterior-posterior MN identities impacted upon the loss of different accessory/insulation factors. The illustrations on the left side of each panel demonstrate the regulation of gene expression through the combinatorial binding of accessory factors in addition to cohesin and CTCF at loop anchors. The right side of each panel shows MN identity determination/transition based on *Hox* gene expression in the absence of various accessory/insulation factors, as compared to WT MNs. (C) Model depicting the homeotic transformations in the skeletal structure of mice. Cervical, thoracic, lumbar, sacral, and caudal vertebra are shown in indicated colors. Stars mark the vertebrae impacted at cervicothoracic and thoracolumbar boundaries in CTCF/MAZ *Hox5|6* binding deficient mice and *Patz1* KO mice, respectively.

## DISCUSSION

The impetus for this investigation was the lack of a credible explanation for the emergence of the varying chromatin boundaries that foster distinctive, critically important changes in the gene expression profile during differentiation. The absence of detectable changes in the genomic occupation of the well-recognized insulator, CTCF, upon differentiation, was incompatible with the dynamic formation of distinctive chromatin boundaries. Given this phenomenon along with the ability of loop-extruding cohesin to travel through chromatin roadblocks larger than its ring size^24,37–40^, we anticipated that a predetermined process for blocking cohesin-mediated loop extrusion during development would involve the participation of specific DNA binding factors that associate with cohesin in a regulated manner. Indeed, we have identified a family of zinc finger proteins, distinct from the CTCF zinc-finger protein, which appear to foster disparate gene expression profiles by establishing appropriate cohesin-associated chromatin boundaries.

Our findings underscore that different chromatin boundaries contain various accessory factors/insulators as identified herein, and that their combinatorial binding with cohesin, with or without CTCF, could function to determine specific gene expression patterns during development. Importantly, skeletal defects observed at the thoracolumbar boundary in *Patz1* KO mice is consistent with the previous studies of *Hox* mutants (*i.e. Hoxc8, Hoxd8, Hox9* and *Hox10*) indicating transitions in thoracic-lumbar regions^41–44^. In addition to their co-localization with cohesin, the impact of the accessory proteins/insulators identified herein on the diversity of the gene expression program during development, is further substantiated by our observation that the ZNFs, MAZ, PATZ1, VEZF1, CTCF, and cohesin exhibit various expression patterns across different tissues or cell types. Moreover, as shown here, CTCF, MAZ, PATZ1, VEZF1, and importantly, the other identified ZNFs all exhibit distinct DNA binding site preferences. Thus, we propose that these identified accessory proteins/insulators function combinatorially at distinct genome sites to promote the formation of discrete, cohesin-mediated chromatin boundaries during differentiation and development.

As the evidence herein supports that lineage-specific expression and/or binding of a subset of these accessory proteins/insulators is determinant to establishing specific boundaries, we conceptualized a model that includes accessory factors (*i.e.* MAZ, PATZ1, ZNF263, and other ZNFs), in addition to CTCF and cohesin (Figure 7). In this model, different combinations of these factors that exhibit tissue-specific expression and specific DNA binding specificity, would dynamically collaborate with cohesin to foster the specific chromatin interactions necessary to establish the appropriate chromatin boundaries. For example, the existence of loops containing PATZ1 binding peaks in regions lacking CTCF binding, and the reduction of looping interactions in the absence of PATZ1 highlight its importance in genome organization. Our model envisions that one or more of the other ZNF members may coordinate with PATZ1 to establish appropriate chromatin boundaries. We speculate that the different permutations of these ZNFs, with the inclusion or lack thereof of CTCF, constitute a significant reservoir of potential genomic events for establishing cohesin-mediated loop formation reflective of different cell lineages. The viability of these permutations would depend on the tissue specific expression of the ZNFs, the presence of their specific DNA-binding sites in the genome, and possibly their interactions. Interestingly, and consistent with our model, cell-type specific distal gene regulation has been shown to be dependent on the cohesin loading cycle through WAPL^45^. Cohesin turnover on chromatin during cellular differentiation could facilitate the *de novo* combinatorial action of cell-specific factors that, upon binding to chromatin, can construct distinct chromatin boundaries^45^.

Notably, previous studies in *Drosophila*^3,4,46,47^ and in mammals^48,49^ report the presence of multiple “insulators” at a given locus and/or their possible homo/hetero-dimerization properties that may increase the strength of insulation. In addition to these scenarios, the binding of such insulators alone or in a combinatorial fashion, as described in *Drosophila,* may provide unique features to local three-dimensional interactions that inherently impact enhancer-promoter functional contacts and consequently gene expression^3^. Indeed, in addition to the known multiple insulator binding proteins in *Drosophila* (*i.e.* BEAF32, dCTCF, Su(Hw), Zw5, GAF, CP190, MOD(MDG4))^4,5^, later reports indicated additional insulators such as Pita and ZIPIC, the human ortholog of which is ZNF276^50,51^. Of note, several insulators such as dCTCF and Su(Hw) in *Drosophila* contain C2H2 zinc-finger domains implicated in DNA-binding and in particular, GAF, MOD(MDG4), and CP190 have BTB/POZ domains^4,52–54^ involved in dimerization^55^ and important for long-range interactions^56,57^, similar to PATZ1 in mammals^58^. Based on our analysis indicating the presence of PATZ1 at loops (Figures 2D-E and S10C-D), the reduction in looping interactions upon loss of PATZ1 (Figure 6 and S9-S10), and the multimerization properties of CTCF in mammals^59^, the BTB/POZ domain of PATZ1 may be critical for homo/heterodimerization during loop formation. Similarly, the SCAN domain within the ZNF proteins (*e.g.* ZNF263) mediates protein-protein interactions^60^, and may facilitate their dimerization. Furthermore, the DNA-binding motifs of MAZ^20^ and PATZ1 (Figure S4A) show similarity to that of CLAMP, which is involved in dosage compensation by promoting chromatin insulation and long-range chromatin interactions in *Drosophila*^61–63^. Hence, we speculate that the candidates identified here through our analysis of the pattern of cohesin re-localization in the absence of previously identified insulators, represent other mammalian insulators/co-factors (Figures 1H and S1E).

This study revealed the roles of PATZ1 and ZNF263 during *in vitro* differentiation of ESCs into MNs and further, the role of PATZ1 during mouse skeletal development. Our findings clarified their impact on cellular identity determination as an outcome of differential chromatin boundary-mediated *Hox* gene expression, which is mechanistically connected to the alterations in looping interactions in the genome. Our findings point to different combinations of the other ZNFs exhibiting key roles in implementing the program whereby various, requisite chromatin boundaries are constructed in a given cell type and during cellular fate determination (Figure 7). As the general approaches described in this study provided the DNA motifs associated with the relocation of cohesin in the absence of CTCF and MAZ, we expect our findings to be broadly applicable to other borders, both *Hox* and non-*Hox*, and to other cellular fates.

### Limitations of the study

Our analysis indicates the possibility of multiple insulation factors that co-localize with cohesin on chromatin. These results are consistent with earlier work in other organisms, and the role of *Patz1* in loop formation has been demonstrated in this study. In addition, the loss of *Patz1* or *Znf263* impacted distinct *Hox* gene borders known to be insulated during differentiation, and the *Hox* gene mis-expression phenotype was observed as skeletal abnormalities in *Patz1* KO mice. However, the function/s of each factor may depend on the cell type, genes impacted, their regulation, underlying chromatin organization, and the overall biological context.

## ACKNOWLEDGEMENTS

We thank L. Vales for reading and advice on the manuscript; Y. Grobler for providing *Drosophila* S2R^+^ cells; D. Hernandez for technical assistance in mice studies; R. Ripert and R. Cruz for further technical assistance. We also thank the NYU Grossman School of Medicine’s Genome Technology Center, particularly A. Heguy, P. Zappile, and P. Meyn, for help with sequencing; the Applied Bioinformatics Laboratories for bioinformatics support; the Rodent Genetic Engineering Laboratory for help with mice generation; and the Cytometry and Cell Sorting Core, particularly P. Lopez and M. Gregory for help with FACS. This study utilized computing resources at the High-Performance Computing Facility of the Center for Health Informatics and Bioinformatics at the NYU Grossman School of Medicine. For the later stages of this work, we thank the Onco-Genomics Shared Resource (OGSR) of the Sylvester Comprehensive Cancer Center at the University of Miami, RRID: SCR_022502, for the help with sequencing.

## Author Contributions

H.O., P.Y.H. and D.R. conceived the project, designed the experiments and wrote the paper. H.O. and P.Y.H performed the experiments and the bioinformatic analysis; E.G.B. helped with mice studies; H.C. performed the initial bioinformatic analysis of ChIP-seq data under the supervision of A.T. and helped with Micro-C intra-TAD analysis; S.Y.K. helped with mouse model generation; and E.O.M. advised on the progression of this study.

## STAR METHODS

### RESOURCE AVAILABILITY

#### Lead contact

Further information and requests for resources and reagents should be directed to and will be fulfilled by the lead contact, Danny Reinberg.

#### Materials availability

The plasmids generated in this study will be deposited to Addgene. All unique/stable materials generated in this study are available from the corresponding author upon reasonable request with a completed Materials Transfer Agreement.

#### Data and code availability

- Sequencing data has been deposited at Gene Expression Omnibus (GEO) and accession (GSE230482) is publicly available as of the date of publication. The list of differentially expressed genes in *Maz* KO mESCs used in Figures S5H and S6D was obtained from the previous study^20^. Publicly available datasets used are listed in Table S1. Raw data of western blots in the figures 1D, 2C, 3B, S5A, and S7C were deposited in Mendeley. The access to raw data in Mendeley is publicly available as of the date of publication at https://doi.org/10.17632/48xw9hxvpm.1.
- This paper does not report original code.
- Any additional information required to reanalyze the data reported in this paper is available from the lead contact upon request.

## EXPERIMENTAL MODEL AND STUDY PARTICIPANT DETAILS

The detailed cell lines and mouse strains/genotypes have been reported in Key Resources Table. This study has been performed under compliance with ethical regulations and approved by NYU/NYULMC Institutional Biosafety Committee. Mouse studies were approved by NYU Grossman School of Medicine’s Institutional Animal Care and Use Committee and University of Miami Institutional Animal Care and Use Committee. Animal housing conditions were as follows: dark/light cycle, 6:30 pm to 6:30 am (off) / 6:30 am to 6:30 pm (on); temperature, 21 °C ± 1 or 2 °C; and humidity range, 30–70%. This work does not report phenotype in relation to the sex of animals.

## METHOD DETAILS

### Cell culture and motor neuron differentiation

E14 mESCs (ES-E14TG2a, ATCC, CRL-1821) were cultured in standard medium supplemented with LIF, and 2i conditions (1 mM MEK1/2 inhibitor (PD0325901, Stemgent) and 3 mM GSK3 inhibitor (CHIR99021, Stemgent)). Cervical MN differentiation was performed as described previously^18,20^. Briefly, mESCs were differentiated into embryoid bodies (EBs) in 2 days, and further patterning was induced with addition of 1 μM all-trans-retinoic acid (RA, MilliporeSigma) and 0.5 μM smoothened agonist (SAG, Calbiochem). For caudalization of MNs, previously described methods were applied^30,31^. mESCs were differentiated into EBs in 2 days, and patterning was induced with 1 μM RA, 0.5 μM SAG, 75 ng/ml mouse WNT3A (R&D systems, Cat. 1324-WN-010/CF) and murine FGF-basic (Peprotech, Cat. 450-33) in a gradient of the described concentrations (50 ng/ml, 100 ng/ml, and 150 ng/ml) for 4 days. Biological replicates stand for independent differentiation experiments performed. 293FT cells (Thermo Fisher Scientific, R70007) were cultured in standard medium as described in the manufacturer’s protocol.

### CRISPR genome editing

Single-guide RNAs (sgRNAs) were designed using CRISPR design tools in Benchling (https://benchling.com). All sgRNAs were cloned into pSpCas9(BB)-2A-GFP (PX458) (Addgene, #48138) or pSpCas9(BB)-2A-Puro (PX459) V2.0 vector (Addgene, #62988). The sgRNAs were transfected into mESCs using Lipofectamine 2000 (Invitrogen), as described previously^18,20^. Single clones from GFP-positive FACS sorted cells or puromycin (InvivoGen)-resistant cells were genotyped and confirmed by sequencing. When necessary, PCR products were further assessed by TOPO cloning and sequencing to distinguish the amplified products of different alleles. The sequencing chromatograms were aligned in Benchling. All sgRNAs and genotyping primers are shown in Table S6.

### Cell line generation

#### *Maz* KO cells in CTCF-AID background

To generate *Maz* KO cell lines in CTCF-AID background, CTCF-AID mESCs^9^ were transfected with sgRNA in PX458 vector targeting the *Maz* locus. Knock-out of *Maz* was confirmed by genotyping, sequencing, and western blot. sgRNA and genotyping primers were described previously^20^.

#### *Patz1* KO cells

To generate *Patz1* KO cell lines, WT E14TG2a mESCs were transfected with sgRNA in PX459 targeting the *Patz1* locus. Knock-out of *Patz1* was confirmed by genotyping, sequencing, and western blot.

#### FH-PATZ1 cells

The mouse *Patz1* coding sequence (NM_019574) with an HA-tag sequence fused to its 5’ end was cloned into a PiggyBac vector (pPB-CAG-3xFLAG-empty-pgk-hph, Addgene, #48754) to create pPB-CAG-3xFLAG-HA-PATZ1-pgk-hph plasmid via Gibson assembly (NEB, #E2611). *Patz1* KO cells were transfected with 50 ng of pPB-CAG-3xFLAG-HA-PATZ1-pgk-hph plasmid and 100 ng of Super PiggyBac Transposase plasmid (System Biosciences, #PB210PA-1) using Lipofectamine 2000 (Thermo Fisher Scientific), and selected with 800 ug/mL of Hygromycin B for stable cells. Expression of 3xFLAG-HA-PATZ1 was evaluated by western blot. FH-PATZ1 cells were used to determine the genomic binding of PATZ1 as the commercially available antibodies were ineffective for ChIP-seq experiments.

#### *Znf263* KO cells

To generate *Znf263* KO cell lines, WT E14TG2a mESCs were transfected with sgRNA in PX459 vector targeting the *Znf263* locus. Knock-out of *Znf263* was confirmed by genotyping, and sequencing.

#### FH-ZNF263 cells

The mouse *Znf263* coding sequence (NM_148924.3) with an HA-tag sequence fused to its 5’ end was cloned into a PiggyBac vector (pPB-CAG-FLAG-HA, a modified version of pPB-CAG-3xFLAG-empty-pgk-hph from Addgene, #48754) to create pPB-CAG-FLAG-HA-ZNF263 plasmid via Gibson assembly (NEB, #E2611). *Znf263* KO cells were transfected with 50 ng of pPB-CAG-FLAG-HA-ZNF263 plasmid and 100 ng of Super PiggyBac Transposase plasmid (System Biosciences, #PB210PA-1) using Lipofectamine 2000 (Thermo Fisher Scientific), and selected with 800 ug/mL of Hygromycin B for stable cells. Expression of FLAG-HA-ZNF263 was evaluated by western blot.

#### Cloning of CβF-PATZ1

To generate CβF-PATZ1, mouse *Patz1* coding sequence was inserted into PCR-linearized CβF vector^20^ via Gibson assembly.

### Immunoprecipitation

For immunoprecipitation from mESCs, nuclear extracts of E14 WT and FH-PATZ1 cells were prepared as described previously^64^. Briefly, buffer A (10 mM HEPES, pH 7.9, 1.5 mM MgCl_2_, 10 mM KC1 and 0.5 mM DTT) was used to remove the cytoplasmic fraction, and buffer C (20 mM HEPES, pH 7.9, 25% (v/v) glycerol, 420 mM NaCl, 1.5 mM MgCl_2_, and 0.2 mM EDTA) was used to extract nuclear proteins. The NaCl concentration in the nuclear extracts was then diluted to 250 mM using BC50 buffer (20 mM HEPES, pH 7.9 and 50 mM NaCl), and a 10% NP40 solution was added to the diluted nuclear extracts yielding a final 0.1% concentration. Anti-FLAG® M2 Magnetic Beads (MilliporeSigma, #M8823) were used for immunoprecipitation, and proteins bound to the beads were released by the addition of SDS loading buffer, resolved on a 10% Bis-Tris gel, and analyzed using the specific antibodies listed in Table S7.

For immunoprecipitation from 293FT cells, CβF-PATZ1 expression plasmids were transfected into 293FT cells using PEI, and nuclei was prepared using TMSD and BA450 buffers, as described previously^65,66^. Briefly, TMSD buffer (20 mM HEPES, 5 mM MgCl_2_, 85.5 g/L sucrose, 25 mM NaCl, and 1 mM DTT) was used for cytosol removal, and nuclear extraction was done in BA450 buffer (20 mM HEPES, 450 mM NaCl, 5% glycerol, and 0.2 mM EDTA). FLAG affinity immunoprecipitation and FLAG peptide elution were performed as previously described^20^.

### Expression analysis

Total RNA was purified from cells with RNAeasy Plus Mini kit (Qiagen) and reverse transcription was performed on 1 μg RNA by using Superscript III (Life Technologies) and random hexamers (Thermo Fisher Scientific). RT-qPCRs were performed in replicates on 100 ng cDNA using PowerUp SYBR Green Master Mix (Thermo Fisher Scientific) on a CFX384 Touch Real-Time PCR detection system (Bio-Rad). The primers are listed in Table S6. For RNA-seq analysis, 1 μg RNA was used to prepare ribominus RNA-seq libraries according to the manufacturer’s protocols by the NYU Genome Technology Center.

### Chromatin immunoprecipitation sequencing

ChIP-seq experiments were performed as described^20,67^. Briefly, cells were fixed with 1% formaldehyde, nuclei were isolated, and chromatin was fragmented to ∼250 bp in size using a Diagenode Bioruptor. ChIP was performed using antibodies listed in Table S7. Chromatin from *Drosophila* (1:100 ratio to mESC-derived chromatin), and *Drosophila*-specific H2Av antibody were used as a spike-in control in each sample. For ChIP-seq, libraries were prepared as described^18,20^ using 1-30 ng of immunoprecipitated DNA.

### Library construction

ChIP-seq libraries were prepared as described^18^. RNA-seq libraries were prepared using KAPA library preparation kits by the NYU Genome Technology Center.

### Micro-C Preparation

The Micro-C library was prepared using the Dovetail® Micro-C Kit according to the manufacturer’s protocol. During the differentiation process, an independent pool of 1 million cells was used for Micro-C experiments in both cellular states in WT versus PATZ1 KO conditions. Briefly, the chromatin was fixed with disuccinimidyl glutarate (DSG) and formaldehyde in the nucleus. The cross-linked chromatin was then digested in situ with micrococcal nuclease (MNase). Following digestion, the cells were lysed with SDS to extract the chromatin fragments and the chromatin fragments were bound to Chromatin Capture Beads. Next, the chromatin ends were repaired and ligated to a biotinylated bridge adapter followed by proximity ligation of adapter-containing ends. After proximity ligation, the crosslinks were reversed, the associated proteins were degraded, and the DNA was purified then converted into a sequencing library using Illumina-compatible adaptors. Biotin-containing fragments were isolated using streptavidin beads prior to PCR amplification. The library was sequenced on an Illumina Novaseq X platform to generate > 800 million 2 x 150 bp read pairs, ensuring sufficient data depth for the loop analysis (see Table S8 for the number of reads in each sample).

### CRISPR-based zygotic injection in mice

*Patz1* KO mice were generated by zygotic injection^68^, as described previously^20,34^. Briefly, 50 ng/μl gRNA template (Synthego) (Table S6) and 100 ng/μl *Cas9* mRNA were injected into ∼150 C57BL/6 embryos and transferred into five pseudopregnant females in the NYU Grossman School of Medicine’s Rodent Genetic Engineering Laboratory. The pups were genotyped by PCR using genotyping primers (Table S6) and Sanger sequencing. Mouse studies were approved by NYU Grossman School of Medicine’s Institutional Animal Care and Use Committee. Animal housing conditions were as follows: dark/light cycle, 6:30 pm to 6:30 am (off) / 6:30 am to 6:30 pm (on); temperature, 21 °C ± 1 or 2 °C; and humidity range, 30–70%.

### Axial skeleton staining with alcian blue–alizarin red

The neonates (postnatal day 0.5) were dissected, and skeletal staining was performed as described previously^20,34^. Embryos were fixed for 4 days in 95% ethanol and stained with Alcian blue stain (0.03% Alcian blue, 80% ethanol and 20% acetic acid) for 24 h with rocking at room temperature.

After the samples were washed with 95% ethanol for 1 h twice, they were transferred to 2% KOH solution for 24 h. The specimens were then stained with Alizarin red solution (0.03% Alizarin red and 1% KOH in water) for 24 h. Finally, the specimens were washed in 1% KOH/20% glycerol for 6 days, 1% KOH/50% glycerol for 10 days and kept in 100% glycerol.

### Data analysis

#### RNA-seq analysis

RNA-seq data was analyzed as described^18^. Briefly, sequence reads were mapped to mm10 reference genome with Bowtie 2 (version 2.3.4.1)^69^ and normalized differential gene expression was obtained with DEseq2 (version 1.26.0)^70,71^. Differential gene expression analysis was performed using the Wald test built into DEseq2 with an FDR cutoff of 0.05. Relevant expression and p-values are listed for differentially expressed genes in Table S2 and Table S4. PANTHER database was used for Gene Ontology (GO) analysis^72^. Relevant GO terms are listed in Table S3 and Table S5. Venn diagrams were drawn by using online tools: GeneVenn (https://www.bioinformatics.org/gvenn/) and Venn Diagram Plotter (https://pnnl-comp-mass-spec.github.io/Venn-Diagram-Plotter/).

#### ChIP-seq analysis

ChIP-seq experiments were analyzed as described previously^20,67^. In brief, sequence reads were mapped to mm10 or dm6 reference genome with Bowtie 2 (version 2.3.4.1) using default parameters^69^. Quality filtering and removal of PCR duplicates were performed by using SAMtools (version 1.9)^73^. MACS (version 1.4.2) was used for narrow peak calling using default parameters of ‘macs2’^74^. After normalization with the spike-in *Drosophila* read counts or total read counts (for RAD21, to avoid normalization artifacts^15^); ChIP-seq heat maps, density plots, and tracks were generated using Easeq^75^ (http://easeq.net<u>, version v 1.111</u>). ChIP-seq read densities were also visualized in Integrative Genomics Viewer (version 2.16.2)^76^. ChIP-seq reads were extended to 150 bp using default parameters in Easeq. ‘ChIPpeakAnno’ package (version 3.20.1) from Bioconductor^77^ was used to draw Venn diagrams to visualize the overlap among ChIP-seq samples. In addition, BEDTools (version 2.27.1) were also used for the assessment of overlaps^78^. Heat maps were also generated using deepTools (version 3.5.4)^79^. ChIP-seq replicates were assessed by visualizing at Integrative Genomics Viewer (version 2.16.2) and generating heat maps. PAVIS tools were used for ChIP-seq peak annotation to the genes^80^. ChIP-seq “bed” file coordinates were converted into “fasta” by using fetch sequences tool within Regulatory Sequence Analysis Tools (RSAT)^81^; MEME (version 5.5.1) was used for motif analysis of PATZ1 in mESCs^82^, and Tomtom (version 5.5.1) was used as a motif comparison tool^83^. Motif search in MEME was performed *de novo* until 1000 sites were reached and corresponding e-values are depicted.

For public data, filtered alignment files from ENCODE (https://www.encodeproject.org/) were directly used for the analysis. When filtered alignment files were not available, raw sequence reads from NCBI were analyzed via the Seq-N-Slide pipeline (https://doi.org/10.5281/zenodo.5550459).

The data used are listed in Table S1. Human reference genome hg38 were used for comparative analysis. ChIP-seq read densities were normalized to fragments per million reads per kbp. Easeq^75^ or pyGenomeTracks^84,85^ was used to generate the heat maps and representative tracks. The UpSet plot was generated using Intervene^86^ (https://asntech.shinyapps.io/intervene/).

#### Analysis of RAD21, CTCF, and PATZ1 occupancy in loop anchors of HepG2 cells and ESCs

Significantly enriched chromatin loops were called using HiCCUPS from the Juicer package^87^ with default parameters. Overlaps between RAD21, CTCF, and PATZ1 ChIP-seq peaks and chromatin loops were analyzed with pairToBed from the BEDTools suite (version 2.27.1)^78^ and pgltools (version 1.2.1)^88^. Visualization of Hi-C and associated ChIP-seq data was made with pyGenomeTracks^84,85^.

#### Micro-C analysis

The Micro-C analysis was performed by the Dovetail® according to the tools described in https://micro-c.readthedocs.io/en/latest/index.html. Briefly, HiC contact matrices were generated in each sample by aligning fastq files and generating valid pairs bam files. Using HiCCUPs from the Juicer package^87^, and Mustache^89^, loops have been identified in each sample with default parameters. Micro-C comparative analysis between WT versus PATZ1 KO samples were performed by the Dovetail® by using HiCcompare algorithm^90^. The standard HiCcompare method applies a joint normalization for WT and KO conditions prior to differential analysis. Thus, the process reinforces the statistical robustness of the loops called. The standard outputs generated include a table of genomic coordinates of pairs of regions detected as differentially interacting, interaction frequencies, the difference, and the corresponding permutation p-value. In addition, an MD normalization plot was generated to visualize the effect of normalization between the conditions to ensure that the data was appropriate for the downstream difference detection (for pipeline, see https://micro-c.readthedocs.io/en/latest/microc_compare.html). AB compartments were generated using FAN-C toolkit^91^ by the Dovetail®, and the results were presented as bar plots in GraphPad Prism.

#### APA analysis, and visualization of loops in ESCs and MNs

Normalized APA analysis was performed for all the loops identified by HiCCUPs in WT versus PATZ1 KO ESCs and MNs using 5kb resolution by using Juicer APA tools^87^. In addition, normalized APA analysis was performed for the common loops between ESCs and MNs upon differentiation and loss of PATZ1 (see Figure S10), using 5kb resolution by using Juicer APA tools^87^. In this analysis, the common loops were obtained by using the loops identified by HiCCUPs and intersecting the loops in WT ESCs and MNs with pgltools (version 1.2.1)^88^. Visualization of the zoomed-in regions in Micro-C was performed using Juicebox^92^. Differential loops were plotted by using HiCcompare results in Rstudio.

## QUANTIFICATION AND STATISTICAL ANALYSIS

Statistical analysis related to experiments have been described above in each section. Statistical analyses of RT-qPCR were analyzed with CFX Maestro Software 2.3 (Bio-Rad).

**KEY RESOURCES TABLE** (Separate word document)

## Supplementary Tables

**Table S1.** Public datasets analyzed in this study, related to Figures 2, S2 and S3

**Table S2.** RNA-seq expression values in WT vs *Patz1* KO mESCs, related to Figure 4

**Table S3.** Gene ontology analysis in WT vs *Patz1* KO mESCs, related to Figure 4

**Table S4.** RNA-seq expression values in WT vs *Patz1* KO MNs, related to Figure 5

**Table S5.** Gene ontology analysis in WT vs *Patz1* KO MNs, related to Figure 5

**Table S6.** DNA oligos used in this study, related to Figures 1-6 and STAR Methods

**Table S7.** Antibodies used in this study, related to Figures 1-5 and STAR Methods

**Table S8.** The number of reads in Micro-C analysis, related to Figures 6, S9-S10, and STAR Methods

## Supplementary Materials

### SUPPLEMENTARY FIGURES

**Figure S1.**
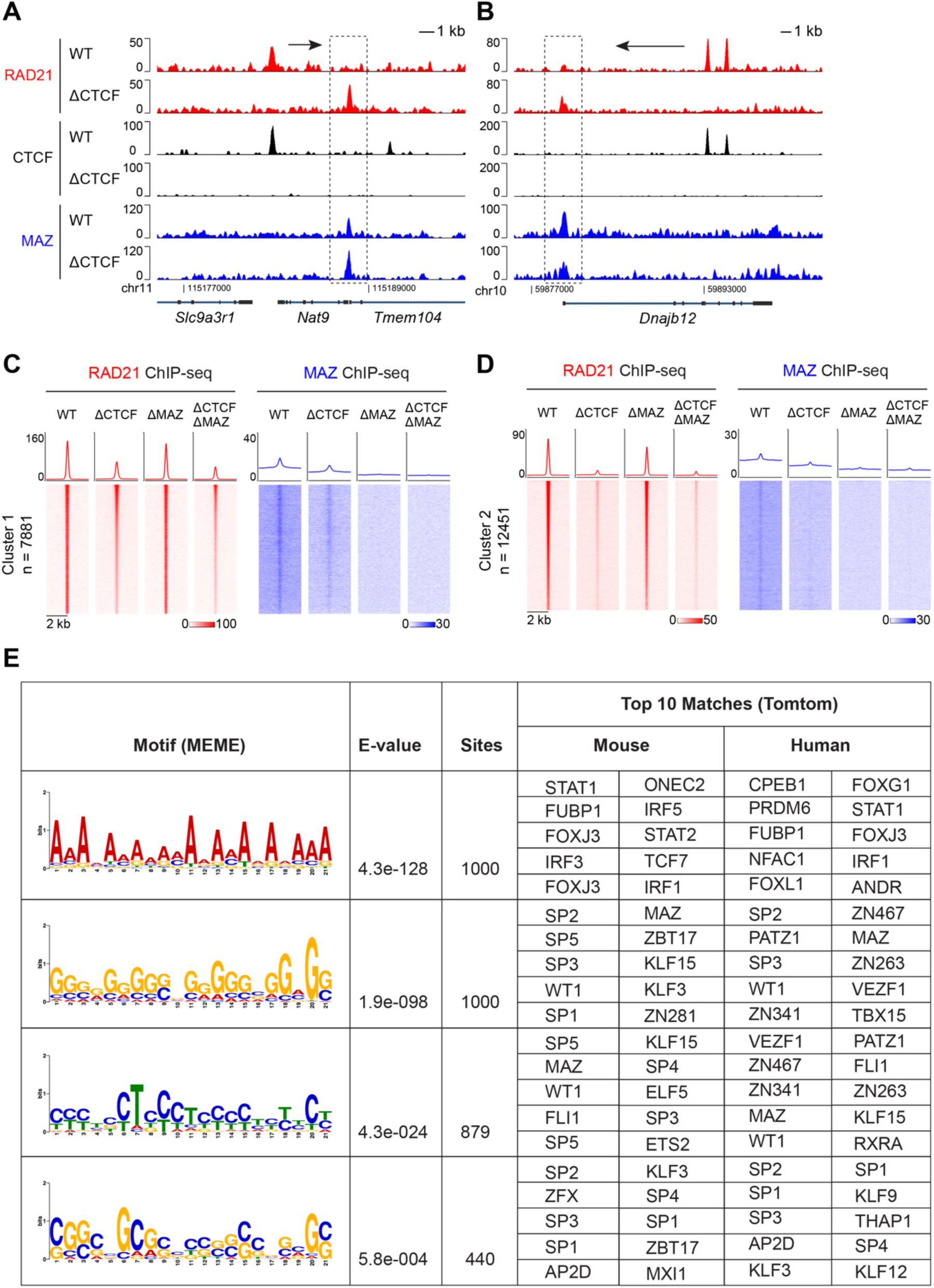
Loss of MAZ in CTCF-degraded mESCs results in the reduction of re-localized RAD21 signal at regions co-occupied by MAZ and RAD21, related to Figure 1. (A-B) Normalized ChIP-seq densities for RAD21, CTCF, and MAZ at (A) *Nat9* and (B) *Dnajb12* loci where re-localized RAD21 overlaps with MAZ binding. ChIP-seq data is from one representative of two biological replicates for CTCF and MAZ, and one biological replicate for RAD21. Additional RAD21 ChIP-seq datasets were reported earlier^15,20^. RAD21-relocalized regions are indicated within dashed-lines. Arrows indicate proximal RAD21 peak to indicate the possible re-localization. (C-D) Heat maps of RAD21 and MAZ ChIP-seq read density in (C) Cluster 1 (n=7,881) and (D) Cluster 2 (n=12,451) within a 4 kb window in WT, ΔCTCF, ΔMAZ, and ΔCTCF/ΔMAZ conditions in ESCs. Average density profiles for ChIP-seq under each condition is indicated above the heat map. ChIP-seq data is from one representative biological replicate for CTCF degron ESCs and one representative of two biological replicates for MAZ KO clones in CTCF degron background. (E) Motif identification for re-localized RAD21 peaks in the absence of CTCF and MAZ by *de novo* MEME motif analysis, along with the corresponding top matches by Tomtom motif comparison tool. Motif search in MEME has been performed *de novo* until 1000 sites were reached and corresponding e-values are depicted in the table.

**Figure S2.**
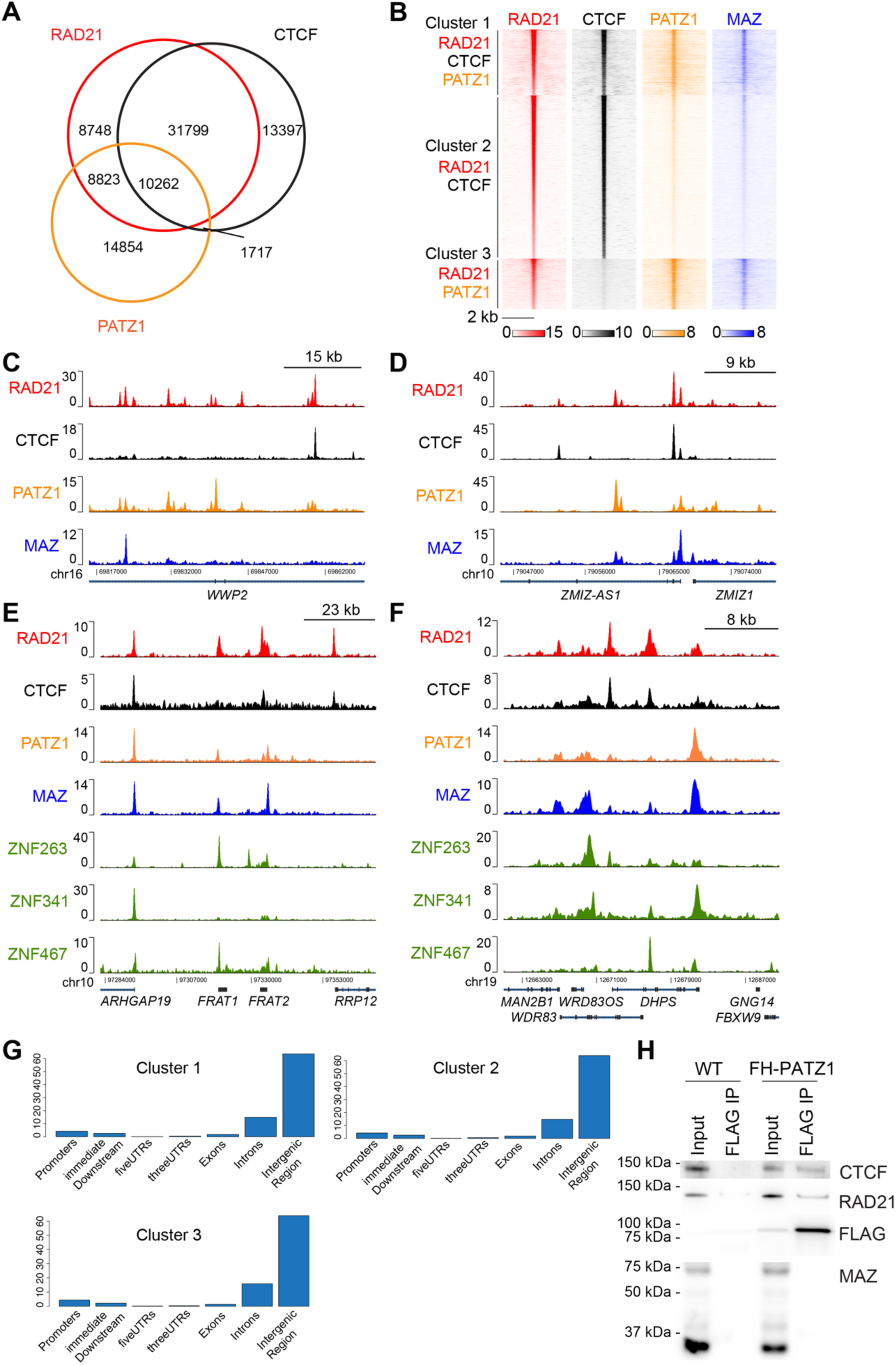
PATZ1 co-localizes with RAD21 on chromatin in HepG2 and HEK293 cells, related to Figure 2. (A) Venn diagram showing RAD21, CTCF, and PATZ1 binding in HepG2 cells. (B) Heat maps of RAD21, CTCF, PATZ1, and MAZ ChIP-seq read density in HepG2 cells clustered as Cluster 1, Cluster 2, and Cluster 3 based on the indicated overlaps with RAD21 signal within a 4 kb window. (C-D) Normalized ChIP-seq densities for RAD21, CTCF, PATZ1, and MAZ, wherein peaks co-localizing with RAD21 and/or CTCF were visualized in HepG2 cells. ChIP-seq data in HepG2 cells is from two combined biological replicates. (E-F) Normalized ChIP-seq densities for RAD21, CTCF, PATZ1, MAZ, and other zinc finger proteins, ZNF263, ZNF341, and ZNF467, wherein RAD21 and/or CTCF co-localizing peaks were observed in HEK293 cells. ChIP-seq data in HEK293 cells is from one replicate for RAD21 and one representative of two biological replicates for others. The source of genomics data used is listed in Table S1. (G) Distribution of RAD21, CTCF, and PATZ1 co-localizing binding sites across genomic features based on Figure 2B. Cluster 1, 2, and 3 refers to the clusters shown in Figure 2B. (H) Western blot analysis of CTCF, RAD21, FLAG, and MAZ upon FLAG-PATZ1 immunoprecipitation from nuclear extract of 293FT cells (n=2, see Figure 2C).

**Figure S3.**
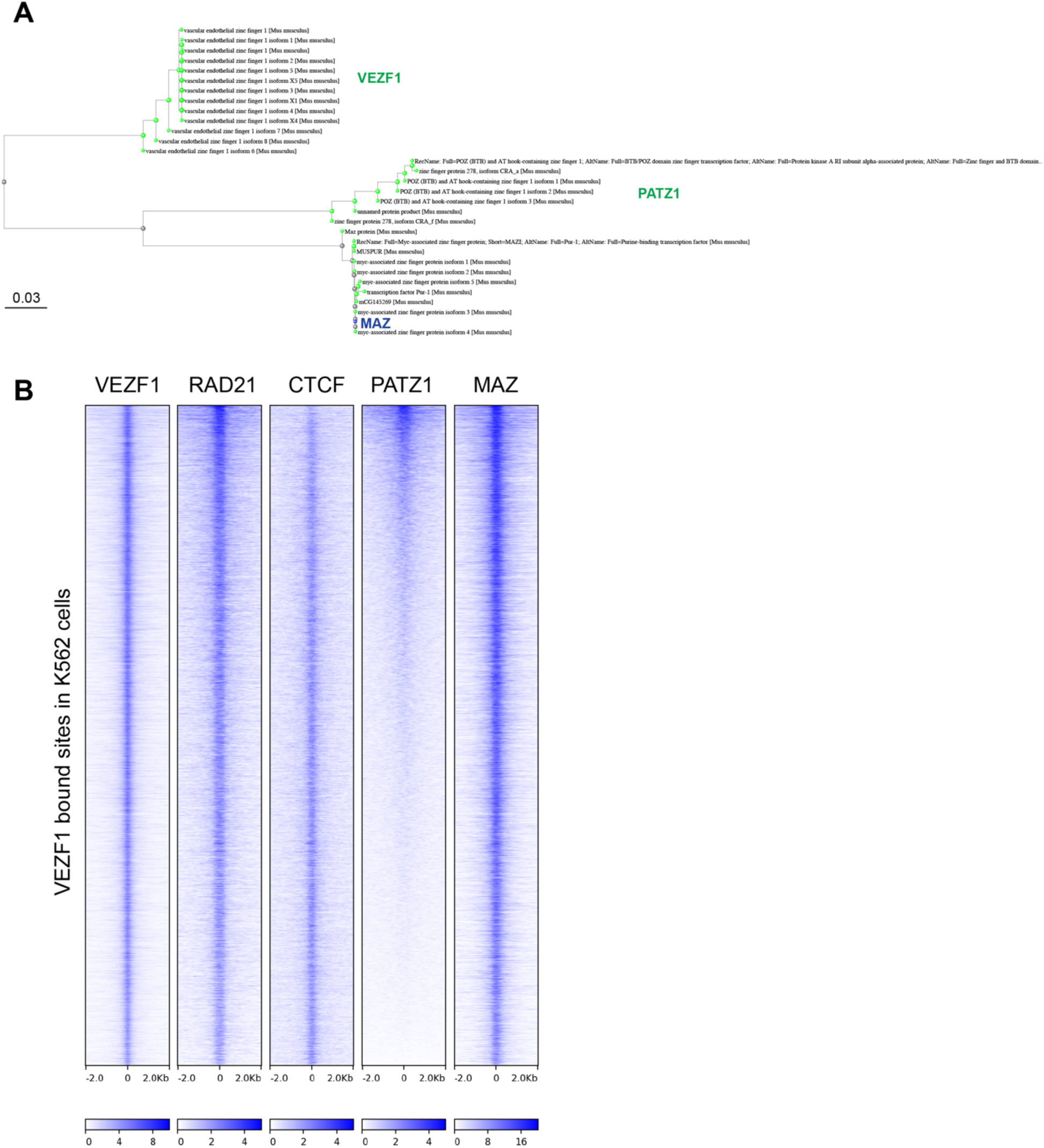
PATZ1 and VEZF1 show protein sequence similarity to MAZ, and co-localize with MAZ, related to Figure 1 and 2. (A) Protein blast (blastp) analysis of MAZ protein. Top 30 matches were drawn as the tree. (B) Heat maps of VEZF1, RAD21, CTCF, PATZ1 and MAZ ChIP-seq read density in VEZF1 binding sites within a 4 kb window in K562 cells. ChIP-seq data in K562 cells is from one representative of two biological replicates. The source of genomics data used is listed in Table S1. VEZF1 ChIP-seq was re-analyzed.

**Figure S4.**
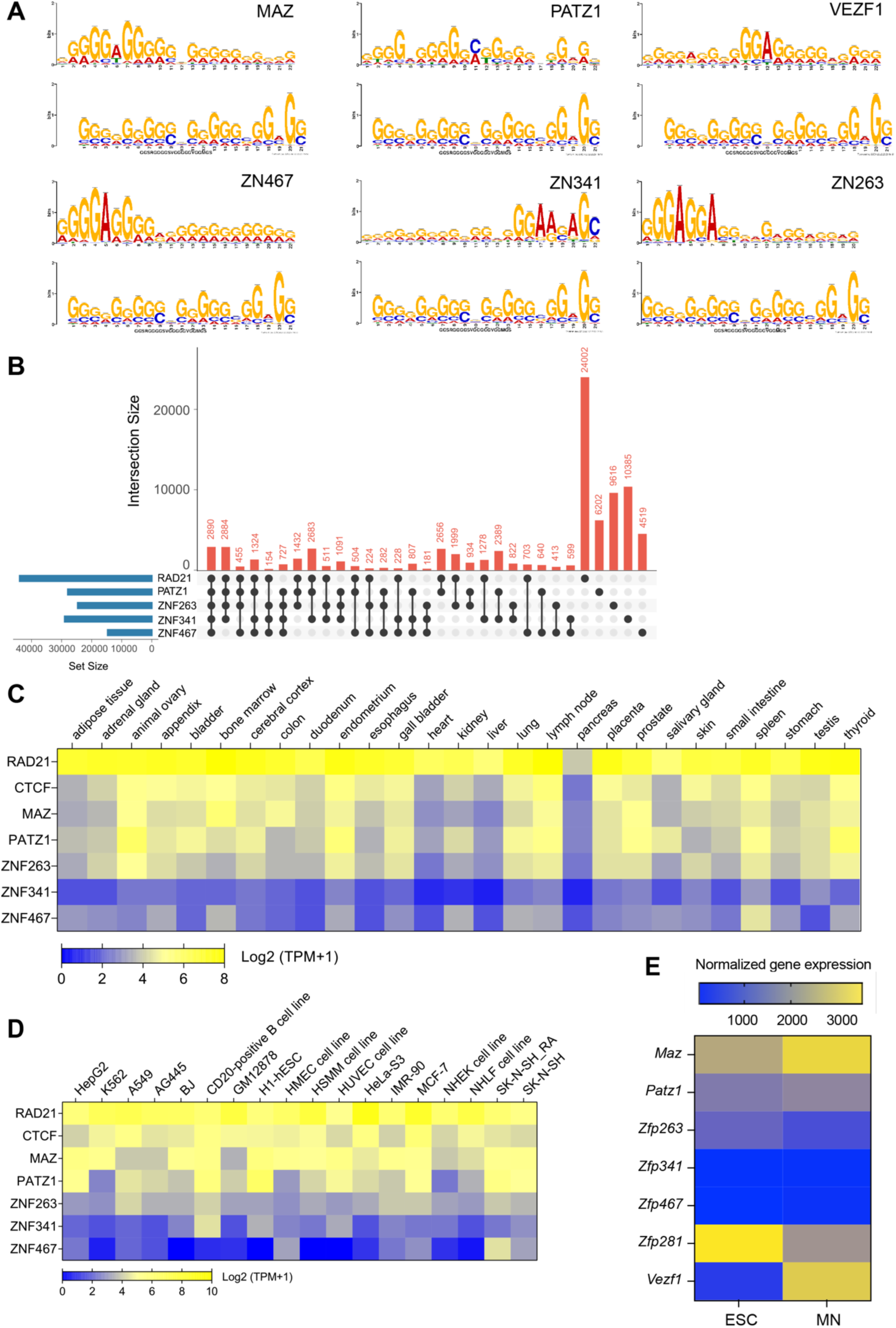
PATZ1 and other ZNFs are expressed at varying levels across different tissues and cell types and co-localize with RAD21 on chromatin, related to Figure 1 and 2. (A) Representative alignment of top motif matches to the candidates: MAZ, PATZ1, VEZF1, ZN467, ZN341, and ZN263. *De-novo* motif analysis was performed as depicted in Figure S1E for re-localized RAD21 peaks in the absence of CTCF and MAZ. Motif alignments were generated through Tomtom motif comparison tool. (B) UpSet plot indicating the overlap of RAD21, PATZ1, ZNF263, ZNF341, and ZNF467 binding in HEK293 cells (see Figure 2F). (C-D) Heat map of RNA-seq expression [log_2_ (TPM+1)] of RAD21 and ZNFs across different human tissues (C) and ENCODE cell lines (D). (E) Heat map of RNA-seq expression (Normalized counts) of MAZ, PATZ1, ZNFs and VEZF1 during ESC to MN differentiation from two biological replicates.

**Figure S5.**
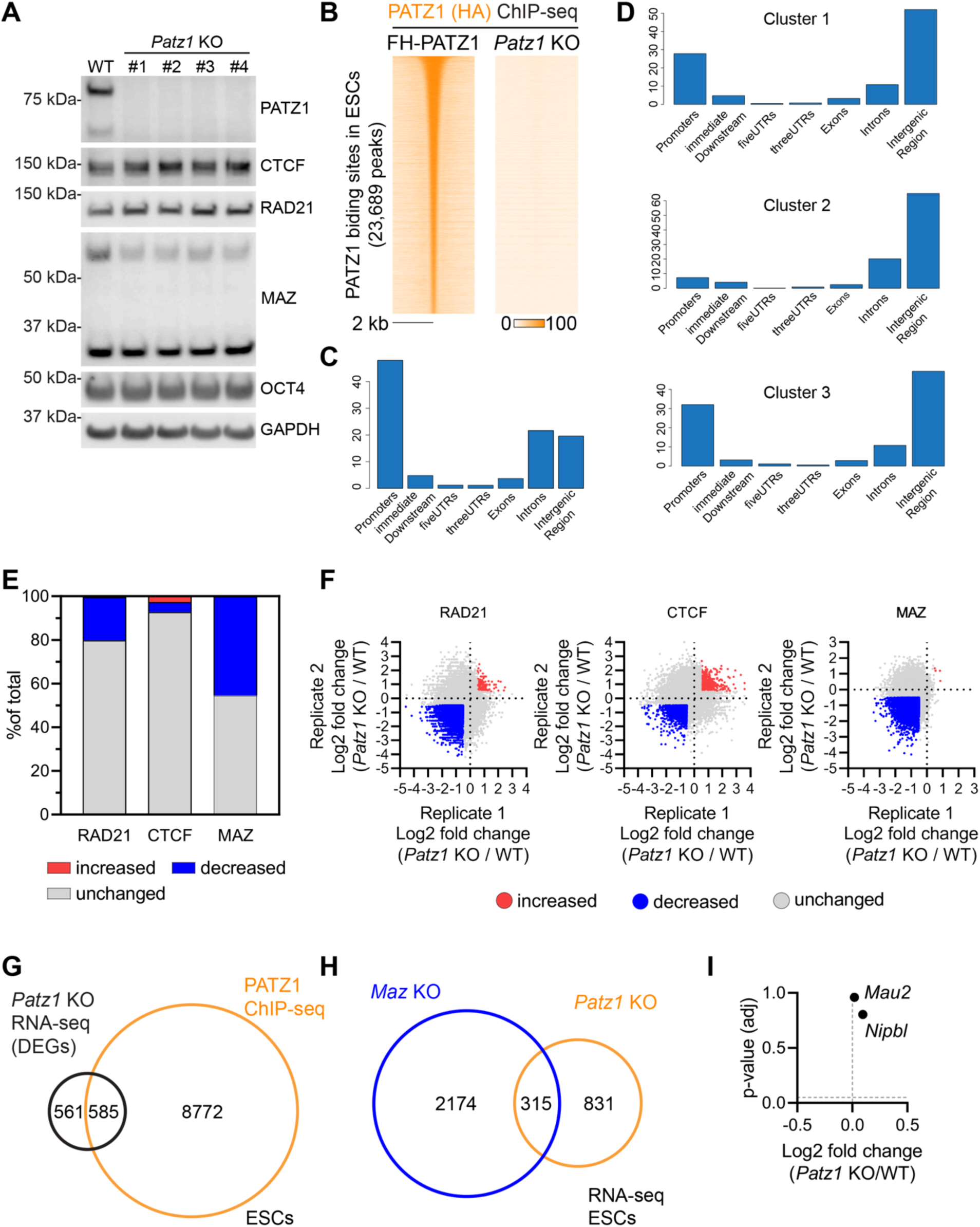
The effects of *Patz1* depletion in mESCs, related to Figures 3 and 4. (A) Western blot analysis of PATZ1, CTCF, RAD21, MAZ, OCT4, and GAPDH in WT and *Patz1* KO clones. (B) Heat maps of PATZ1 (HA) ChIP-seq read densities in FH-PATZ1 and *Patz1* KO mESCs at PATZ1 peaks, showing the specificity of PATZ1 ChIP-seq signals. (C) Distribution of PATZ1 binding sites in ESCs across genomic features. (D) Distribution of RAD21, CTCF, and PATZ1 co-localizing binding sites across genomic features based on Figure 3D. Cluster 1, 2, and 3 refers to the clusters shown in Figure 3D. (E) Changes in ChIP-seq densities of RAD21, CTCF, and MAZ in *Patz1* KO with the cutoff of ±0.5 log_2_(fold change) observed in two independent biological replicates. (F) Ratio of RAD21, CTCF, and MAZ ChIP-seq densities between *Patz1* KO and WT from 2 independent biological replicates. (G) Overlap of PATZ1 ChIP-seq signal with the differentially expressed genes (DEGs) in *Patz1* KO ESCs compared to WT ESCs. (H) Venn diagram indicating the overlap of differentially expressed genes upon *Patz1* KO and *Maz* KO in mESCs from three biological replicates. (I) The fold change in expression of the cohesin loader genes, *Nipbl* and *Mau2*, in mESCs was assessed by RNA-seq from three biological replicates.

**Figure S6.**
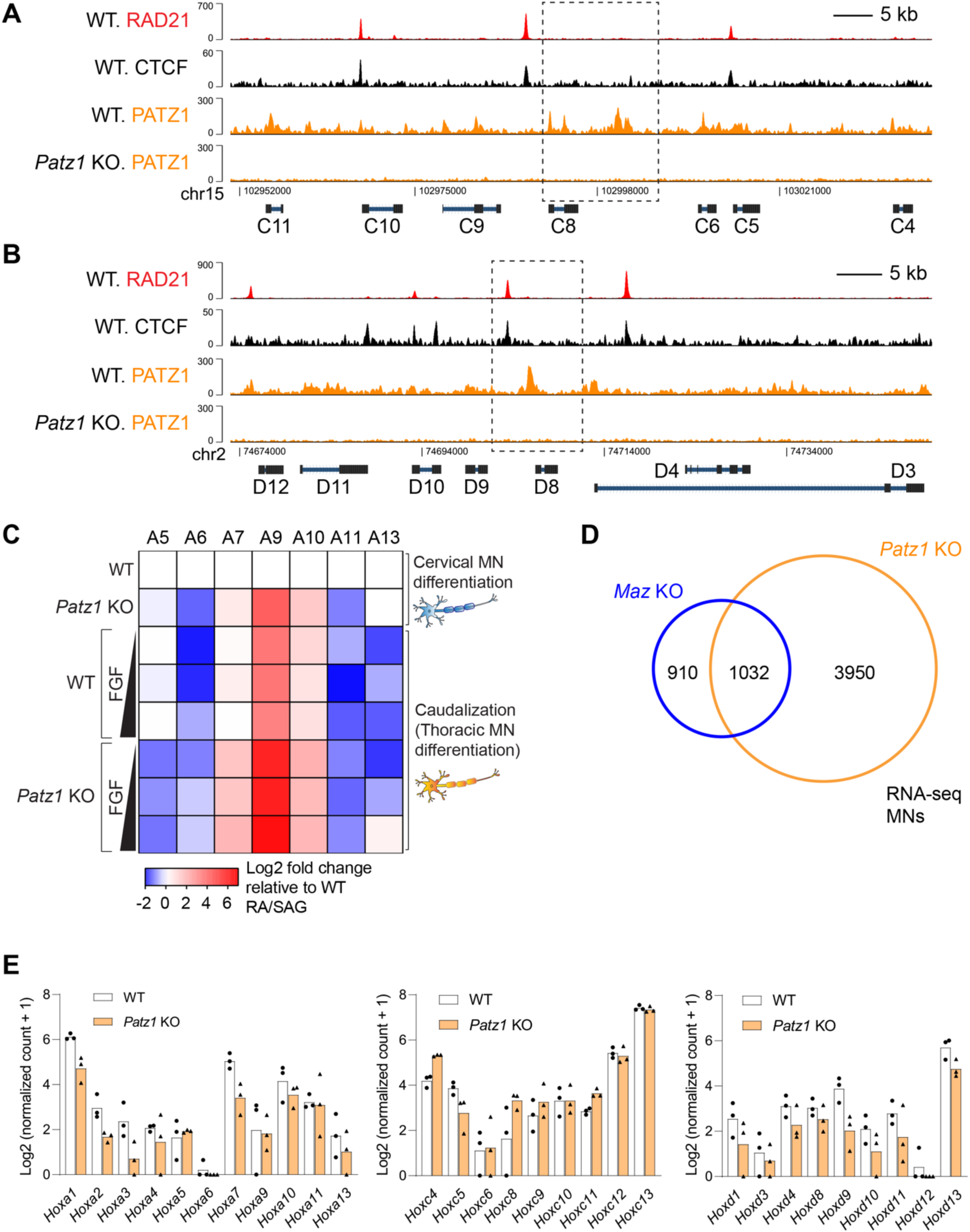
Loss of PATZ1 results in de-repression of *Hoxa9*, *Hoxc9* and *Hoxd9* in cervical MNs, indicating a rostro-caudal patterning defect in MNs, related to Figure 5. (A-B) Normalized ChIP-seq densities for RAD21, CTCF, and PATZ1 in WT and *Patz1* KO mESCs at the indicated region in the *HoxC* (A) and *HoxD* (B) cluster. (C) Heat map of relative gene expression in WT versus *Patz1* KO MNs at the *HoxA* cluster in cervical MNs and thoracic MNs from one biological replicate. WNT/FGF signaling gradient was utilized for MN caudalization, as described previously (see Methods for details). (D) Venn diagram indicating the overlap of differentially expressed genes upon *Patz1* KO and *Maz* KO in motor neurons from two and three biological replicates, respectively. (E) RNA-seq normalized counts for *Hox* gene expression across *HoxA, C,* and *D* clusters in WT versus *Patz1* KO ESCs from three biological replicates.

**Figure S7.**
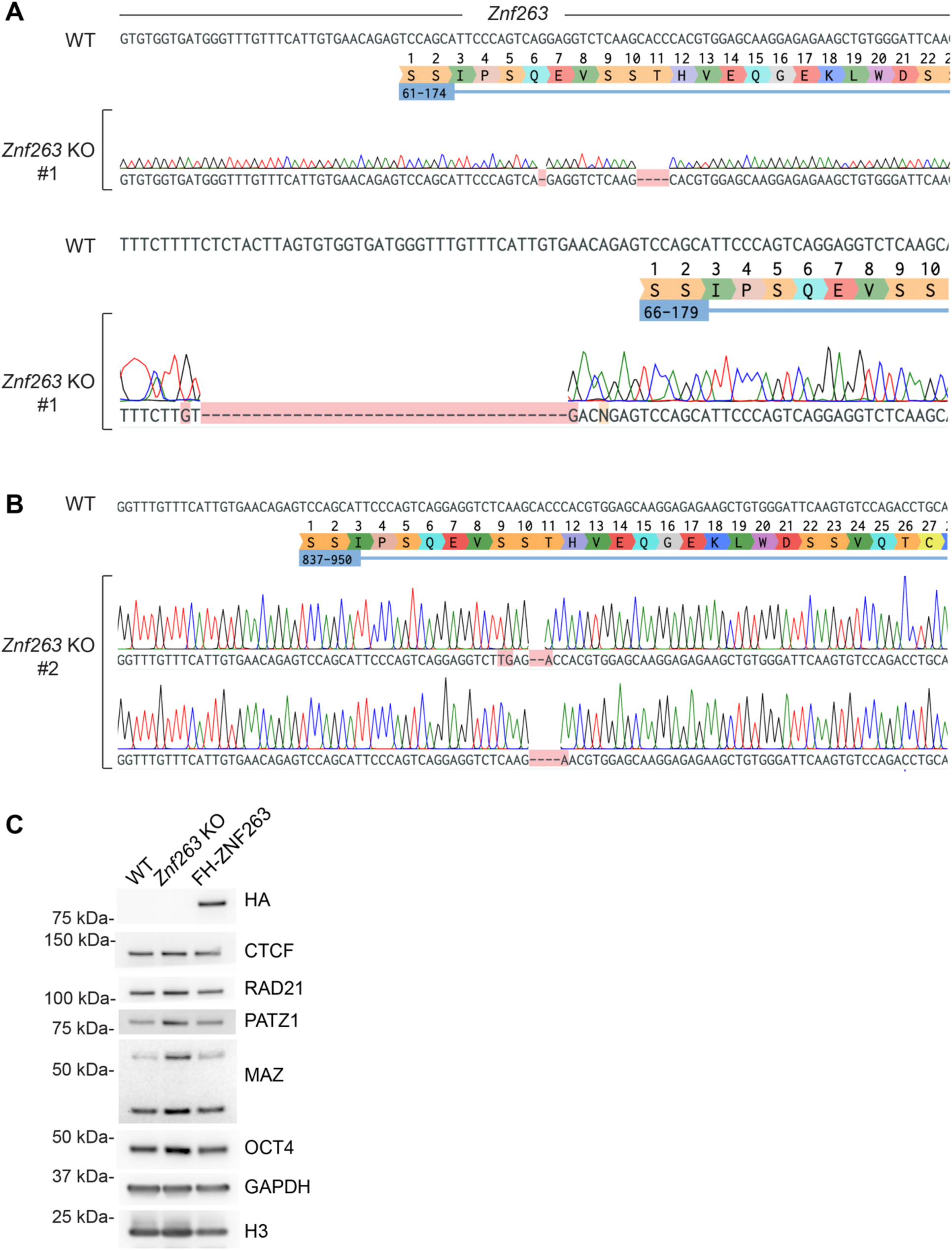
*Znf263* KO mESC generation via CRISPR and FH-ZNF263 expression in mESCs, related to Figure 5. (A-B) CRISPR based deletions in two mESC clones upon targeting of the *Znf263* locus (see Figure 5). Clone #1 (A) and clone #2 (B) harbor the indicated deletions resulting in frame-shift mutations. (C) Western blot analysis of FH-ZNF263 (HA), CTCF, RAD21, PATZ1, MAZ, OCT4, GAPDH, and Histone H3 in WT, *Znf263* KO, and FH-ZNF263 mESCs.

**Figure S8.**
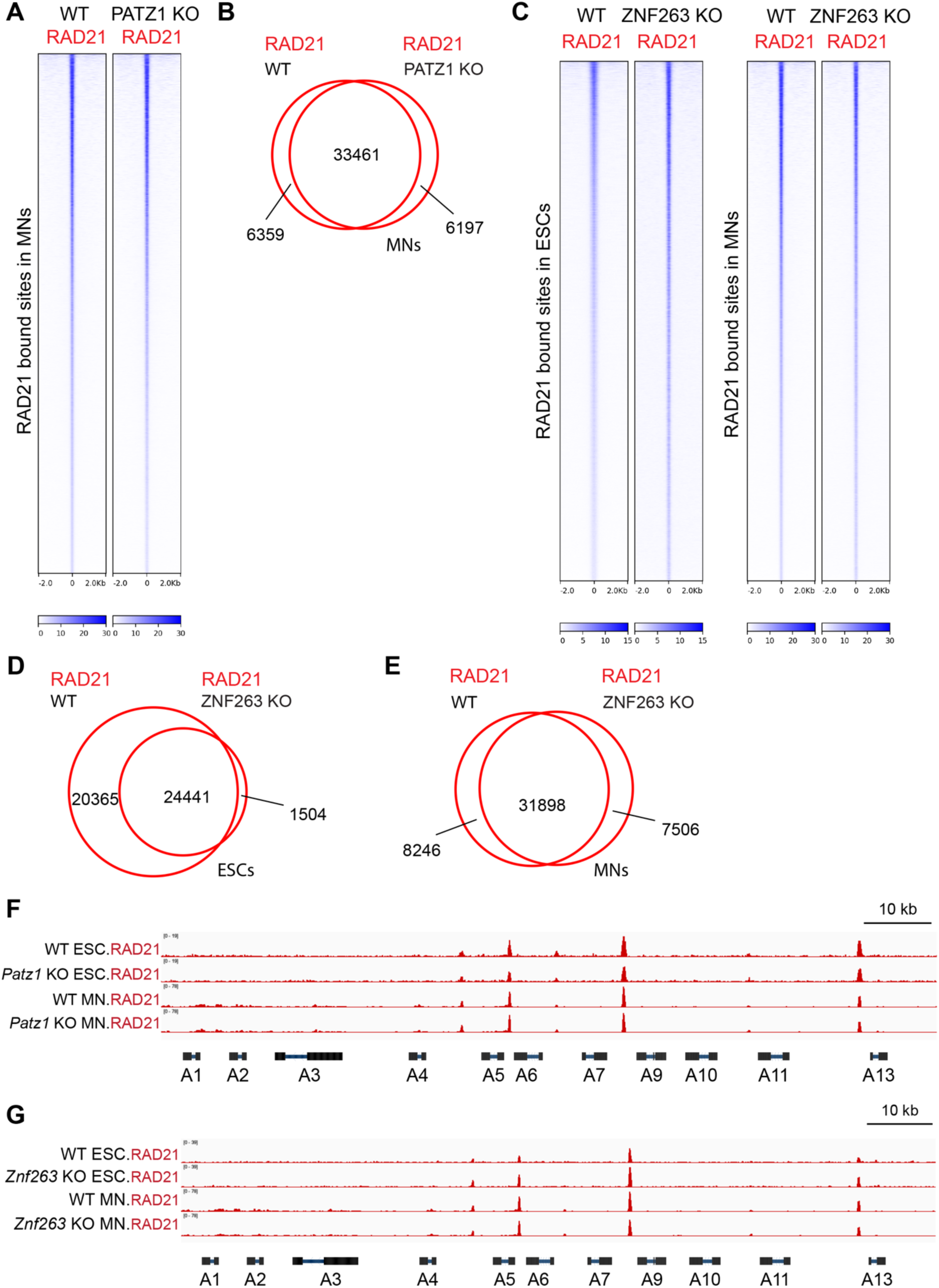
RAD21 binding on chromatin is impacted upon loss of PATZ1 or ZNF263, related to Figure 4 and 5. (A) Heat maps of RAD21 ChIP-seq read density in RAD21 binding sites within a 4 kb window in WT vs PATZ1 KO MNs. (B) Venn diagram indicating the overlap of RAD21 binding sites in WT vs PATZ1 KO MNs. (C) Heat maps of RAD21 ChIP-seq read density in RAD21 binding sites within a 4 kb window in WT vs ZNF263 KO mESCs and MNs. (D) Venn diagram indicating the overlap of RAD21 binding sites in WT vs ZNF263 KO mESCs. (E) Venn diagram indicating the overlap of RAD21 binding sites in WT vs ZNF263 KO MNs. (F) Normalized ChIP-seq densities for RAD21 in WT versus *Patz1* KO in mESCs and MNs at the *HoxA* cluster. (G) Normalized ChIP-seq densities for RAD21 in WT versus *Znf263* KO mESCs and MNs at the *HoxA* cluster.

**Figure S9.**
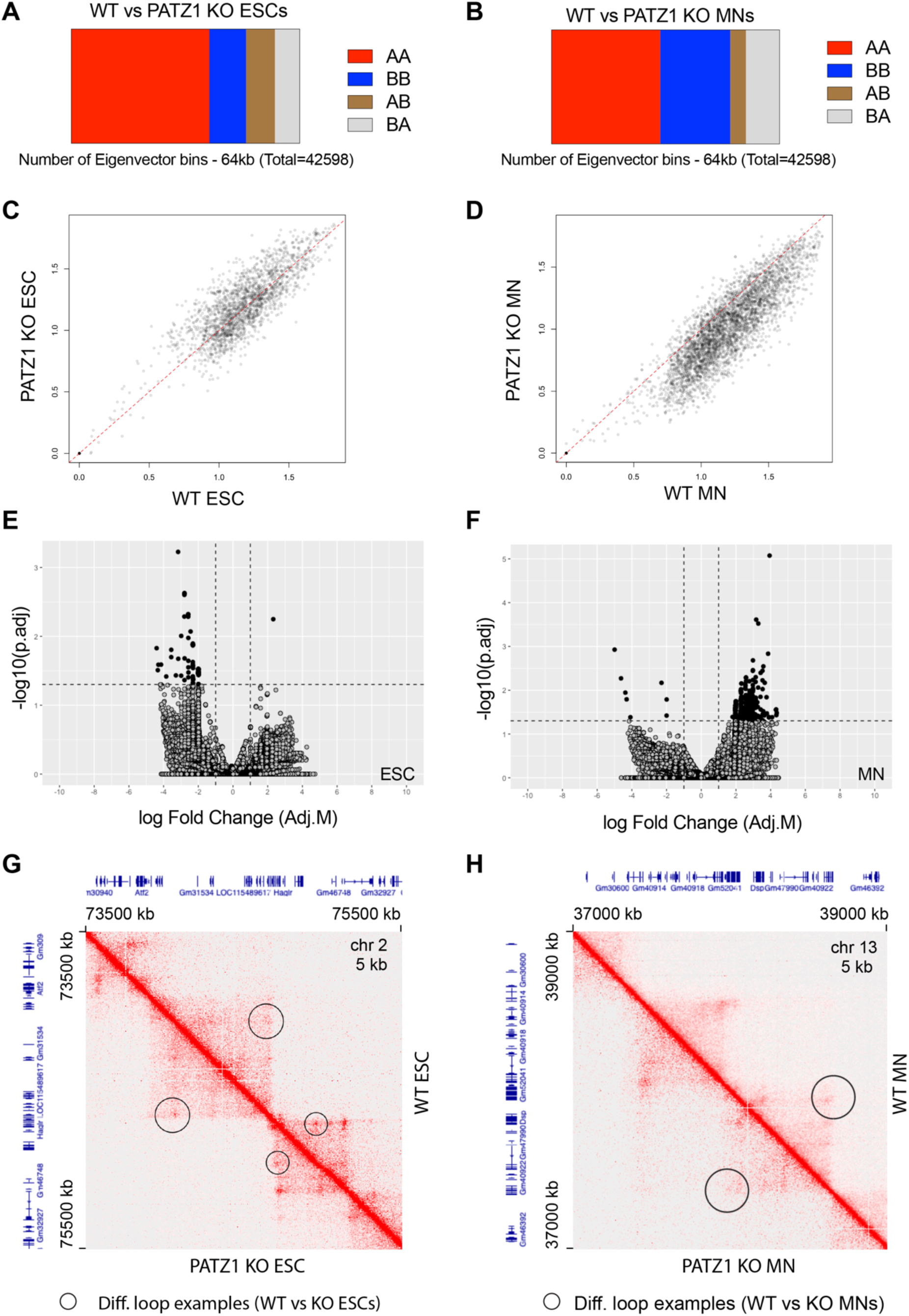
Loss of PATZ1 results in changes in the looping interactions at the level of genome organization in ESCs and MNs, related to Figure 6. (A-B) Bar plot showing AB compartments in WT versus PATZ1 KO ESCs (A) and MNs (B). (C-D) Scatter plot showing intra-TAD activity scores in WT versus PATZ1 KO ESCs (C) and MNs (D). The scores indicating intra-TAD activity from arrowHead (10 kb) has been plotted in WT versus PATZ1 KO. (E-F) Volcano plot of differential loop analysis in WT versus PATZ1 KO ESCs (E) and MNs (F). Black circles indicate the values with p.adj < 0.05, and gray circles indicate all loops included in the comparative analysis from WT and KO conditions via HiCcompare algorithm (see STAR methods). (G-H) Visualization of Micro-C contact matrices for a zoomed-in region (G) around the *HoxD* cluster in WT versus PATZ1 KO ESCs, and (H) the *Dsp* locus in WT versus PATZ1 KO MNs. Examples of the differential loops detected are indicated with black circles. The resolution is 5 kb. Shown above and left are gene annotations.

**Figure S10.**
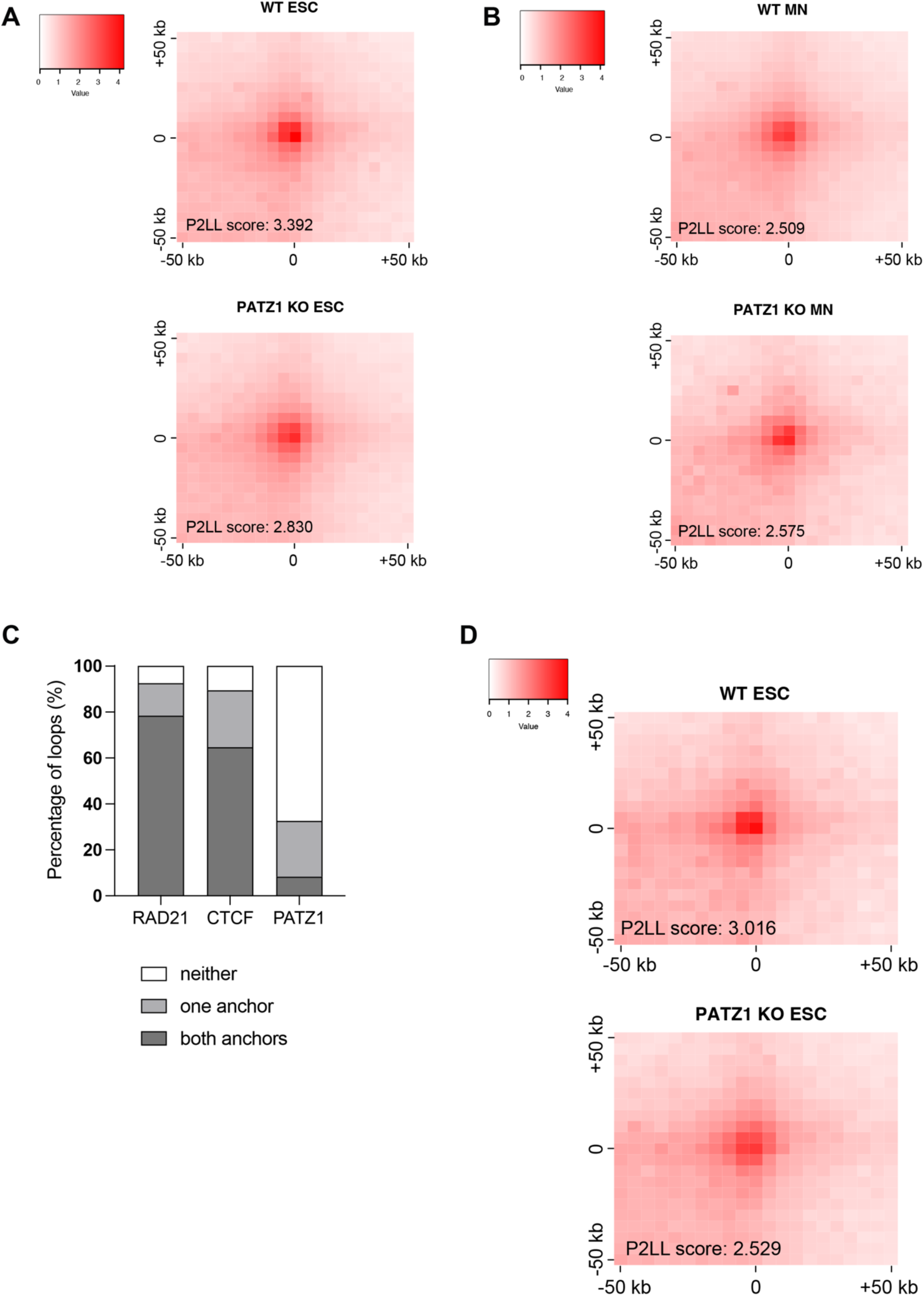
Loss of PATZ1 results in alterations in looping interactions at common loops in ESCs and MNs, and PATZ1-occupied regions in ESCs, related to Figure 6. (A-B) Normalized APA plots of all common loops between ESCs and MNs plotted in WT versus PATZ1 KO (A) ESCs and (B) MNs. The loops common between WT ESCs and WT MNs have been generated using pgltools as described in STAR methods. The resolution of APA is 5 kb. P2LL (Peak to Lower Left) is the ratio of the central pixel to the mean of the mean of the pixels in the lower left corner (see Table S8 for Micro-C sequencing reads). (C) Percentage of Micro-C loops in ESCs overlapping with RAD21, CTCF, and PATZ1 ChIP-seq peaks. (D) Normalized APA plots of loops in WT versus PATZ1 KO ESCs at PATZ1 ChIP-seq co-occupied regions. The resolution of APA is 5 kb. P2LL (Peak to Lower Left) is the ratio of the central pixel to the mean of the mean of the pixels in the lower left corner (see Table S8 for Micro-C sequencing reads).

**Figure S11.**
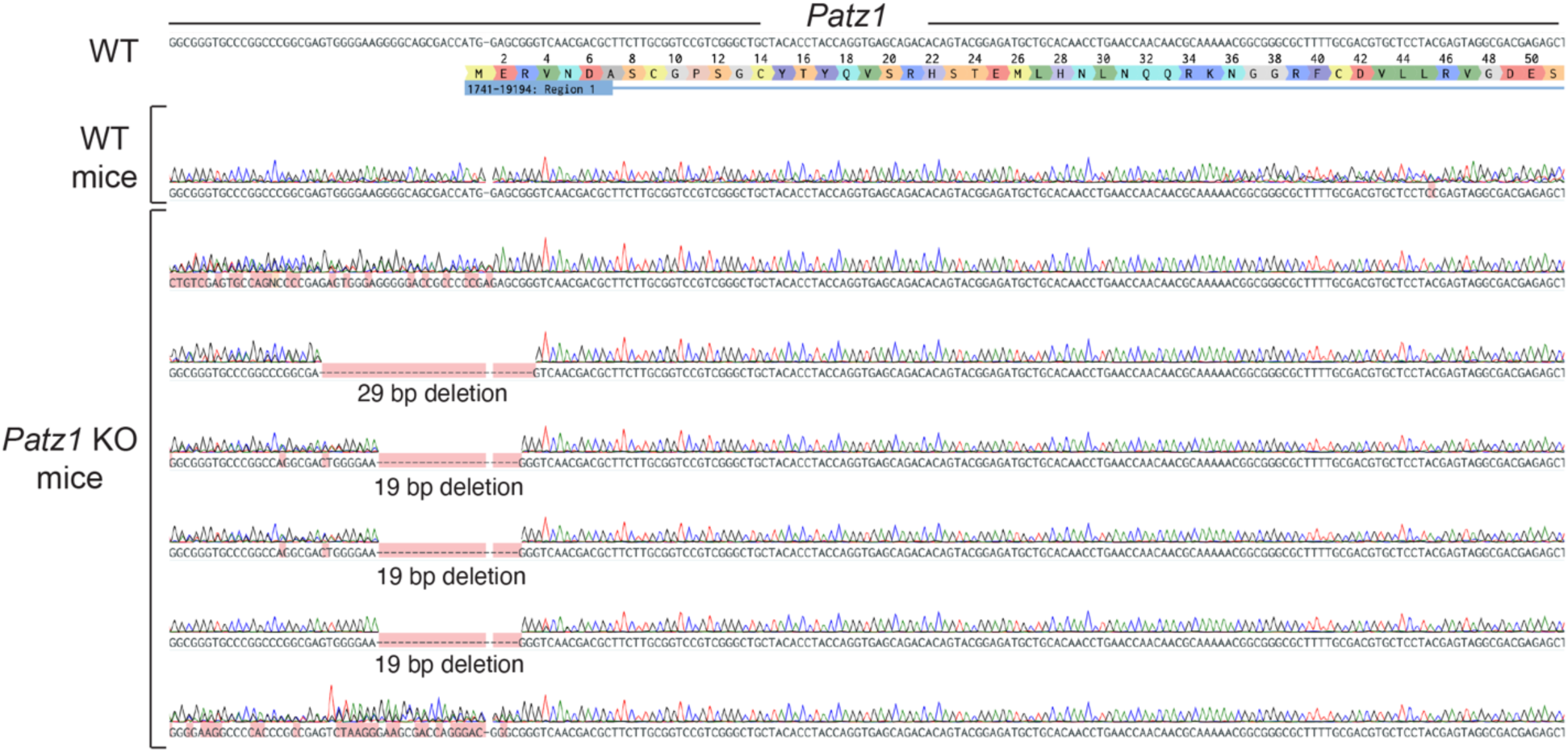
*Patz1* KO mice generation via CRISPR-based zygotic injection *in vivo*, related to Figure 6. CRISPR based deletions in 6 mice upon targeting of the *Patz1* locus (see Figure 6G). 4 mice harbor the indicated deletions resulting in frame-shift mutations, while two mice appear to have heterozygous alleles.

